# Salience-based information integration: An overarching function of the “social brain”?

**DOI:** 10.1101/2023.01.30.525877

**Authors:** Claire Lugrin, Arkady Konovalov, Christian Ruff

## Abstract

Behavior in social contexts is routinely accompanied by neural activity in a brain network comprising the bilateral temporoparietal junction (TPJ), dorsomedial and dorsolateral prefrontal cortex (dmPFC and dlPFC), and precuneus. This network – often referred to as the “social brain network” (SBN) – is thought to have evolved in response to the information processing demands of life in social groups. However, its precise functional contributions to behavior are unclear, since many of its areas are also activated in non-social contexts requiring, for example, attentional orienting or context updating. Here we argue that these results may reflect a basic neural mechanism implemented by areas in this network that is commonly required in both social and non-social contexts: Integrating multiple sensory and memory inputs into salient configurations, such as social constellations or perceptual Gestalts. We tested this hypothesis using a numeracy paradigm that orthogonally varied the salience of sensory target configurations and the required motor responses. Even in this non-social task, several regions of the SBN (TPJ, dmPFC, and precuneus) showed higher activity when the goal required the brain to attend to more versus less salient perceptual configurations. This activation pattern was specific to configuration salience and did not reflect general task demand or switching to new contexts. Taken together, these results suggest that the integration of information into salient configurations may be a key function of SBN regions, thus offering a new perspective on the widespread recruitment of these areas across social and non-social contexts.

## INTRODUCTION

The term “social brain network” (SBN) refers to a set of cortical regions commonly associated with a wide range of social behaviors, including the bilateral temporoparietal junction (TPJ), dorsomedial and dorsolateral prefrontal cortex (dmPFC and dlPFC), and precuneus^1,2^. According to a prominent hypothesis^3^, these regions may have evolved to process the complex information required to navigate social groups^4^. Support for this hypothesis comes from findings that neocortex size correlates with the number of individuals in primate groups^3^, the complexity of the social interactions between individuals in these groups^5,6^, and the social network size in humans^7,8^ and rhesus macaques^9^. This relationship seems to be specific to the complexity of the social environment, since neocortex size is not related to non-social aspects of the environment associated with different cognitive demands (e.g. size of habitat)^3^.

Based on these considerations, various theories have proposed that the SBN is involved in specific aspects of social cognition and social information processing^1,2,10,11^. For example, the dorsomedial prefrontal cortex (dmPFC) and temporoparietal junction (TPJ, and in particular the right TPJ) have been consistently associated with “theory of mind” (TOM) - the capacity to understand other’s mental states^12–16^ as well as other social processes, such as social learning^17–19^, or empathy^10,20–24^. The dorsolateral prefrontal cortex (dlPFC) is also recruited during social interactions requiring a variety of norm-related, morality and strategic behaviors^25–29^. Pathological changes in the functioning of the SBN are thought to underlie brain disorders such as autism spectrum disorder (ASD), which is associated with malfunctioning social behavior^30^, in particular with difficulties engaging in theory of mind^31^ or perspective taking^32^.

However, in addition to consistent recruitment in contexts requiring social cognition, many SBN regions are also activated in purely non-social contexts. For instance, the right TPJ has also been identified as a key node of the attention reorientation network^33^ and routinely responds to behaviorally relevant non-social stimuli across multiple modalities^34–36^. Some studies have defined different anatomical or functional subregions within the rTPJ that may support different functions^37–39^, but meta-analyses^40,41^ and within-subjects measures across domains^42–44^ have revealed a degree of overlap between activations of the rTPJ in social versus non-social contexts. Moreover, the bilateral TPJ, precuneus, and dmPFC have also been described as the “default mode” network^45^ (DMN), which is commonly activated at rest and deactivated during cognitively demanding tasks. Finally, prefrontal areas described as part of the SBN have also been found to be more generally involved in cognitive control^46,47^ and more general behavioral flexibility^48–50^.

Leaving aside the question of whether these overlaps in fMRI activations during social and non-social contexts indeed reflects the involvement of the same neuronal sub-populations, the large variety of domains associated with activity in the SBN areas thus raises several questions (see also^51,52^ for an in-depth discussion). What computations could these regions implement? Is there a shared functional mechanism that is necessary for social information processing and other types of behavior^53^? What specific aspects of social cognition could rely on this mechanism?

Theoretical accounts of the functional contributions of the SBN areas – and the right TPJ in particular - vary from proposing highly specialized^41,54^ to domain-general^55,56^ roles. The rTPJ is part of the association cortices and receives input from many areas^37,41,57^, processing information from different senses^58,59^. Its neuroanatomical links with multiple networks make it a good candidate for the role of an integration hub, processing signals of different origins, from memory to perceptual and attentional signals^38,60–62^. According to specialized accounts, this anatomical setup of the rTPJ predisposes it to integration mechanisms that are selectively related to the perception and evaluation of social stimuli^38^. For example, the Nexus model argues that this region is responsible for constructing social contexts^41^, while the theory of mind hypothesis proposes that a subregion of the rTPJ is specialized in representing other people’s thoughts^12,13,54^.

Domain-general theories, on the other hand, propose that the integration performed by the rTPJ serves to guide behavior across multiple domains, for instance as a crucial part of the attention reorientation network^55^ that acts as a filter to prevent reorienting attention to all types of environmental stimuli^33,63^, or as a region dedicated to updating internal models of the environment when new sensory information changes expectations about upcoming events^56^. However, the rTPJ is also recruited in the absence of environmental changes and is more broadly associated with multisensory integration of environmental stimuli^60,61,64^. All these theories thus converge in suggesting that the TPJ, and other SBN regions in the association cortex, are key candidates for information integration; but each theory focuses on a particular type of task requiring such integration (i.e., TOM, attention reorientation, contextual update). Here, we build on these existing theories and propose a higher-level role of these regions that could unify the different approaches.

Our proposal is centered around the idea that efficient control of behavior necessitates internal representation of world states, as well as the actions that are best suited for behavioral success in these states (Figure 1d). These representations of action-relevant world states need to be high-level in the sense that they must extract the action-relevant gist from the many inputs the brain processes at any moment of time, such as sensory information, interoceptive signals, and memories (Figure 1d). Thus, the brain needs to contain mechanisms that integrate all these diverse inputs and group them into higher-level conceptual representations (or configurations) that have the right level of abstraction for efficient behavioral control. Gestalt examples are good illustrations of this integration process, where multiple stimuli are grouped in a specific way that constitutes a salient *configuration^65^* of elements. We propose that due to their anatomical layout that integrates connections from many neural systems, the SBN regions may be specialized for such integration of information into high-level configurations.

**Figure 1.**
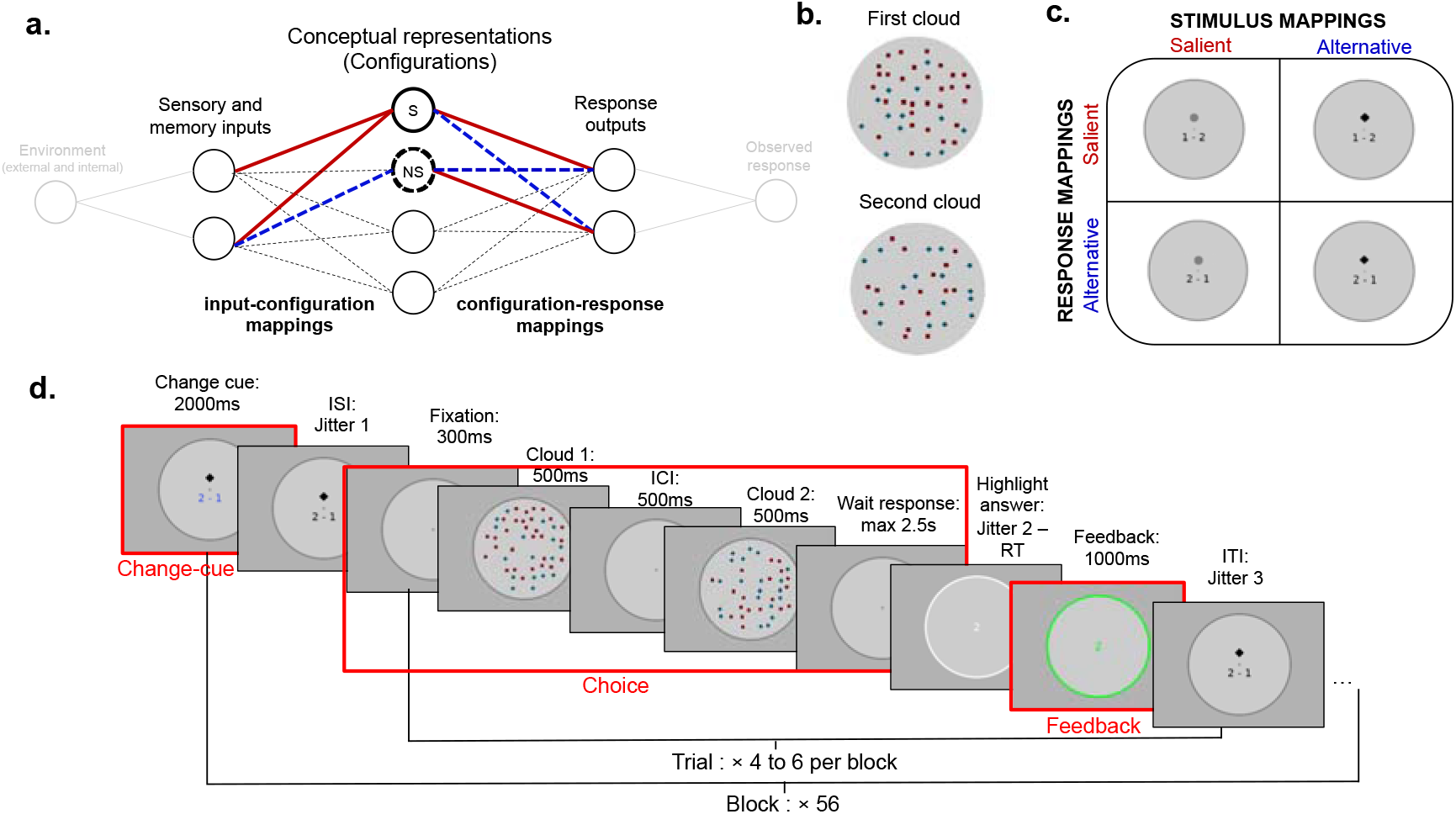
Experimental design and theoretical framework. (**a**) Theoretical framework. A schematic choice model can be decomposed in two parts. In the first part (input-configuration mapping) inputs from the environment are integrated to construct conceptual representations of world states. In the second part (configuration-response mappings) these representations are mapped onto the appropriate motor response. Inputs can be mapped onto salient configurations (S) in an automatic, habitual way (salient mappings indicated in red). Other mappings reflect more atypical and effortful behaviors (NS, alternative mappings indicated in blue). We hypothesize that differential BOLD activity in the SBN reflects the type of input-configuration mapping and does not correspond to changes in the configuration-response mapping. (**b**) Example stimuli. Participants view two consecutive clouds composed of two types of elements (squares and diamonds). The difference in the number of elements between the two clouds, as well as the difference in the number of alternative stimulus elements (squares or diamonds) systematically varies. (**c**) 2X2 task design. The subjects faced four types of trials (stimulus vs response, salient vs alternative mappings). In the salient stimulus mapping (Sal Stim) conditions the subjects had to select the cloud containing the largest total number of elements. In the alternative stimulus mapping (Alt Stim) conditions, they had to select the cloud with the largest number of specific elements (e.g., diamonds). In the salient response mapping (Sal Resp) conditions the subjects had to press a button once to select the first cloud, and twice to select the second cloud; in the alternative response (Alt Resp) mapping conditions this correspondence was reversed (**d**) fMRI task design. The subjects compared the two clouds and reported the cloud containing the most elements. They played 56 blocks of 4 to 6 trials. At the beginning of each block, a cue indicated the change of one condition (stimulus or response mapping). The subjects then observed two consecutive clouds and selected the cloud containing the most relevant elements. Response was followed by choice feedback (1 or 2), followed by accuracy feedback (green for correct, red for incorrect). The events included in the fMRI analysis are indicated by a red rectangle.

In particular, we propose that these areas should be more involved when input-configuration mappings are *“salient”*, i.e., when they correspond to a prepotent, habitual way of grouping information. For example, when we see “:-)”, our brain readily maps this onto the configuration of a smiling face. Similarly, when we see a movie scene where a male person is on his knee presenting a box to a standing female person, we immediately piece together the different cues (gestures, box, facial expressions) and infer that we are witnessing a marriage proposal. However, the same inputs could also be interpreted in an *alternative* way when the behavioral goals are different. Instead of processing all inputs at once, our brain can focus on particular pieces of information and construct alternative representations. “:-)” could be viewed as three meaningless punctuation marks when the goal is to count the number of characters on a page, and the proposal scene could be analyzed in terms of clothes and fashion accessories if the aim is to judge in which decade the movie was shot. Typically, such alternative input-configuration mappings are less intuitive and can be more effortful and challenging; they may nevertheless be required in some situations depending on the context or behavioral goal.

Based on these considerations and the previous literature, we thus propose that activity in many SBN regions - most notably the TPJ and prefrontal areas - could systematically relate to the difference in processing between such salient and alternative mappings. This hypothesis would explain the increased recruitment of these regions in integrating information from multiple streams^38,41^ and in Gestalt perspectives^65^. It could also explain the consistent involvement of these regions in social behaviors such as TOM^10,12,16,40^, since attributing intentions to others requires grouping multiple behavioral cues (such as context, gesture, facial expressions, etc.) and is something most individuals automatically do when interacting with peers or viewing social situations (especially when the experimental task explicitly demands such attribution). In fact, individuals attribute social intentions even to sets of moving geometric shapes^66^, which suggests that such salience-based integration into behaviorally relevant constellations happens easily and automatically.

Interestingly, individuals with autism spectrum disorder do not automatically attribute social intentions or personality to the geometrical shapes in such tasks^67^. ASD is associated with disorders of social behavior^30,31,32^, but is also accompanied by disorders in non-social processes, such as perceptual information integration^68^. Individuals with ASD often focus on local rather than global information processing; they do not always perceive Gestalt representations as whole salient objects^69,70^ but rather view the different elements of a configuration separately. Functional deficits of the SBN, which are common for ASD individuals^71–73^, could thus be linked to the weaker representation of salient configurations in both social and non-social settings.

To test our proposal, we designed a non-social numeracy paradigm (Figure 1) that varied the characteristics of the input-configuration integration, using identical sensory stimuli but varying the task from focusing on salient mappings (salient representation of the stimuli) to alternative mappings (a non-intuitive representation). We purposefully used non-social stimuli to identify domain-general functional mechanisms that are clearly not specific to social contexts, so that none of our findings could reflect truly social-specific cognition such as mentalizing or updates in other social beliefs.

To rule out potentially confounding factors, we designed our task so that we could control for two alternative accounts of why activity in SBN regions may differ between salient and alternative mappings. First, we controlled for the effect of task demand on this network’s response. We did so because activity in the DMN^45^ - which overlaps substantially with the SBN^74^ - deactivates with increasing task demand in a variety of tasks^45,75,76^. Responses to salient and alternative stimuli mappings may indeed differ in response speed and accuracy, which are common measures of task demand. Since our account proposes a more specific function of the SBN regions that reflects the way information is integrated, rather than general task demand, we controlled for task difficulty by constructing parallel salient and alternative mappings in the response domain that matched the difficulty of the salient and alternative stimulus mappings. Second, since previous research suggested a role of the rTPJ in contextual updating^56^, we also reasoned that our design needs to explicitly control for changes in the stimuli and responses mappings, indicated by contextual cues. We thus also varied the type of contextual switches (between stimulus or response mappings) between blocks of trials and studied how they affected behavior and brain activity, independently of the salience-based integration demands inherent to the current stimulus and response instructions.

## RESULTS

### Sensory and motor salience are reflected in behavior

To test our hypothesis that the SBN regions selectively respond to the salience of input-configuration mappings, independently of potential responses to general task demand or contextual updates, our experimental task systematically varied the salience of both input and response mappings, as well as the types of perceptual and motor switches (Figure 1). The salience of input-configuration and configuration-response mappings was manipulated by means of two stimulus integration and two response rules (Figure 1c). Subjects observed two consecutive clouds of square- and diamond-shaped elements (Figure 1b, see Methods for details) and had to indicate which of the two clouds contained more elements. In the salient stimulus (Sal Stim) mapping condition, subjects had to indicate which cloud had the largest total number of elements (which was, by design, the most salient representation of the cloud). In the alternative stimulus (Alt Stim) mapping condition, the subjects had to indicate which cloud had the largest number of specific elements (squares or diamonds, a subset of the cloud). Thus, this task required the subjects to integrate a subset of the stimuli into a non-salient configuration (Figure 1c) that was less automatically perceived.

As salient and alternative stimulus mappings introduce differences in task demand, we controlled for these differences by designing a Salient and an Alternative response mappings that were matched in difficulty to the salient and alternative stimulus mappings. In the salient response (Sal Resp) mapping condition, subjects responded by pressing the button once to choose the first cloud and twice to choose the second cloud. In the alternative response (Alt Resp) mapping condition, we reversed this mapping (Figure 1c). This task was inspired by the Stroop^77^ and Simon^78^ effects, in which habitual associations between stimuli and response patterns influence subjects’ accuracy in giving non-habitual responses. Thus, these response mappings allowed us to measure how general differences in task demands affect activity in SBN areas, independently of the specific demands associated with salience-based integration of stimuli into configurations.

Subjects switched between the two input-configuration or the two configuration-response mappings every few trials (Figure 1d). Fifty-four subjects played 280 trials of the task (70 per condition) while we recorded their brain activity using 3T fMRI. We first confirmed that salient and alternative stimulus and response mappings were indeed perceived more and less automatically, respectively, by comparing subjects’ accuracy and response time (RT) across conditions. Both these indicators showed that subjects responded to our saliency manipulations (Figure 2a).

**Figure 2.**
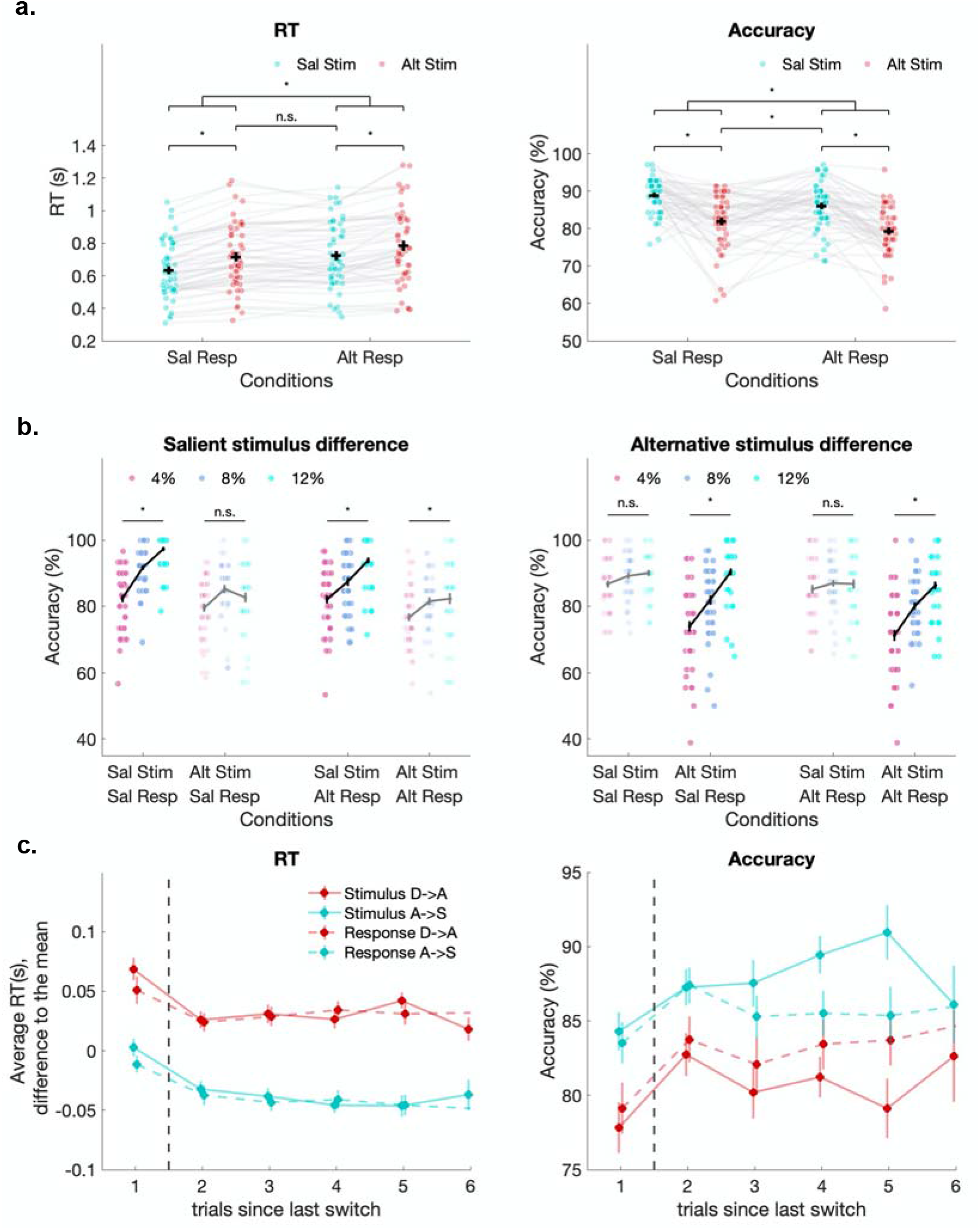
Behavioral data confirm the suitability of our design for studying salience effects in stimulus and motor mappings. (**a**) Average response time (RT) and accuracy in the 4 experimental conditions. Task difficulty, reflected by accuracy and RT increases for the alternative compared to salient mappings in both stimulus and response domains. Each dot and grey line represent one subject, error bars represent s. e.m. The upper lines and stars represent significance (p < 0.025) of paired t-tests between conditions. (**b**) Accuracy for the three levels of stimulus discriminability of each stimulus mapping. The 4%, 8%, and 12% stimulus differences correspond to differences in the proportion of elements between the two clouds. In the salient stimulus plot (left), the difference is computed between the total number of elements. In the alternative stimulus plots (right) the difference is computed between the number of Alt Stim elements (squares or diamonds). Each dot represents one subject, error bars represent s.e.m. The upper lines and stars represent significance (p < 0.05) of linear regressions between the accuracy and discriminability levels. Discriminability, as a second measure of stimulus salience, has a selective effect on subjects’ behavior when the task goal and stimulus which salience is manipulated align. (**c**) Evolution of response time (RT) and accuracy throughout blocks of trials of the same condition. The mean RT is subtracted for each subject, the difference to the mean is represented. The first trial immediately follows a switch, whereas trials 2 to 6 correspond to stay trials. Stimulus switches are represented in plain lines, response switches in dashed lines. The colors represent the direction of the switch: the switches from salient to alternative mappings are represented in red, the switches from alternative to salient mappings are in blue. Error bars represent s. e. m. Stimulus and response switches elicit similar switch costs (RT and accuracy differences between switch and stay trials). As expected, switching to alternative mappings is more costly than switching to salient mappings.

Compared with the condition in which both stimulus and response mappings are salient (Sal Stim/Sal Resp: accuracy = 89%, RT = 628 ms), subjects were significantly less accurate and slower to respond in all alternative conditions (Alternative stimulus (Alt Stim/Sal Resp): accuracy 82%, RT = 711 ms; Alternative response (Alt Resp/Sal Stim): accuracy = 86%, RT = 718 ms; Alternative stimulus and response (Alt Stim/Alt Resp): accuracy = 79%, RT = 784 ms). Mixed effect models with random intercepts and slopes for each subject (Supplementary Table 1) confirmed the significance of the main effects on both accuracy (stimulus type p < 0.001, response type p < 0.001, Supplementary Table 1a) and RTs (stimulus type p < 0.001, response type p < 0.001, Supplementary Table 1b). Interactions between the stimulus and response type did not significantly affect accuracy (p = 0.45, Supplementary Table 1a) and had a small but significant effect on RT (p < 0.001, Supplementary Table 1b). Participants were slightly faster in the Alt Stim/Alt Resp condition than what would be expected with independent effects of the alternative mappings. This difference was however very small and primarily indicated that the difficulty of the two alternatives did not increase compared to each alternative taken separately. Most importantly, accuracy decreased, and RT increased in the alternative stimulus compared to the salient stimulus condition, for both salient (Accuracy: - 6.9%; RT: + 83ms) and alternative responses (Accuracy: −6.7%; RT: +64 ms), showing that the salience in the stimulus and response mappings were independent. Thus, these results validate our experimental design and confirm that we did create well-matched salient and alternative mappings in both response and stimulus domains.

Our task provided a second measure of configuration’s salience, or difficulty of sensory input integration: stimulus discriminability. Within each condition, we systematically varied the relative proportion of attended elements between the two clouds, using differences of 4%, 8%, and 12%. This discriminability of the relevant stimulus (whole cloud for the Sal Stim condition, Alt Stim elements for the Alt Stim condition) significantly increased subjects’ accuracy (Figure 2b, Supplementary Table 1a: mixed effect regression p < 0.001). This measure thus provides a second way of studying the brain activity linked to salient stimulus integration.

To study the alternative hypothesis that SBN regions respond to contextual updating^56^, we analyzed different types of context updates on subject’s behavior and brain activity. We studied the effect of switching from one mapping to the other by comparing trials immediately following a switch (switch trials) with trials occurring later in a block (stay trials). Subjects were less accurate and slower for switch compared with stay trials (Figure 2c). Mixed effect models (Supplementary Table 1) confirmed the significance of the effect of the trial position within a block on both accuracy and RT (Supplementary Table 1, mixed effects, accuracy p = 0.002, RT p < 0.001) confirming the effect of switching costs on behavior. We manipulated the type of task switches (stimulus or response, towards salient or towards alternative mappings). The cost of switching in the stimulus and response domain was matched: the decrease in accuracy and increase in RT following stimulus and response switches was matched for both switch directions (salient to alternative Sal -> Alt and alternative to salient Alt -> Sal, Figure 2c), allowing us to later compare the brain activity for switches of identical costs.

To study the brain’s responses to the different mappings, we created a set of diverse trials by varying additional task parameters (such as the total number of elements or the congruence between the salient and alternative stimulus conditions) in a counterbalanced manner (see Supplementary Methods). Accuracy and reaction time did not significantly depend on these task parameters (Supplementary Figure 1 a-b). Our experimental design therefore allowed us to specifically study the effects of the salience of stimulus and response mappings without other difficulty confounds.

### Social brain regions respond selectively to salient stimulus configurations

Having established the effects of stimulus and motor salience on behavior, we tested whether these effects were indeed differentially reflected in brain activity, using fMRI data.

First, we validated the hypothesis that the SBN selectively responds to the salience of stimulus mappings. Using a linear model (GLM1, see Methods), we contrasted the BOLD signal between the salient and alternative stimulus mappings (Sal Stim > Alt Stim). We found higher activity in the bilateral TPJ, dmPFC, and precuneus in the Sal Stim > Alt Stim contrast (Figure 3a, Supplementary Table 2). These clusters overlapped with the meta-analysis clusters obtained from Neurosynth^79^ using the term “social” (See Supplementary Table 3 for overlap extent). The left anterior dlPFC, which has also been associated with social computations (although absent in the selected meta-analysis) was also significantly more active in the Sal Stim condition. These results support the hypothesis that SBN regions are recruited when stimuli are integrated into salient configurations.

**Figure 3.**
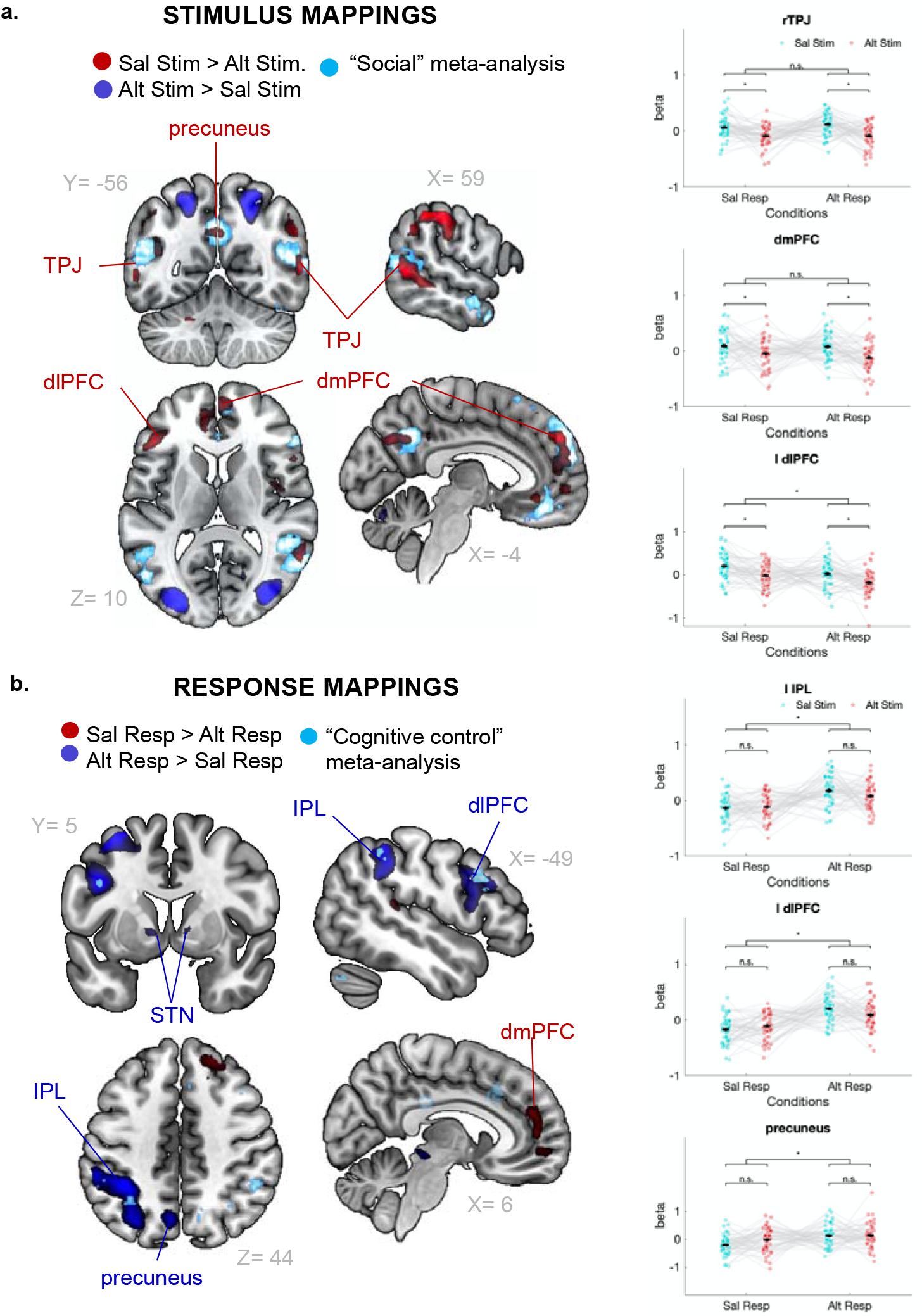
SBN areas respond selectively to salient input-configuration mappings. **a**) Choice event, salient versus alternative stimulus contrast (Sal Stim > Alt Stim). Clusters with significantly higher activity for salient > alternative stimulu-sconfiguration conditions (in red) or higher activity for alternative > salient stimulus-configuration conditions (in blue). The cyan overlay represents ROIs for the term “social” identified by meta-analysis. The demeaned regression coefficients (betas) of GLM4 for the main ROIs are displayed for each subject (dots and grey lines). The upper lines and stars represent significance (p < 0.025) of paired t-tests between conditions using the main contrasts ROIs. These statistical comparisons are different from the main effect contrast used to define the ROIs and are shown for illustration purposes. They show that rTPJ and dmPFC do not respond to differences in response mappings, while the dlPFC ROI is less active the alternative response as well as stimulus condition. (**b**) Choice event, salient versus alternative response contrast (Sal Resp > Alt Resp). Clusters with significantly higher activity for salient > alternative (in red) and alternative > salient configuration-response conditions (in blue). The cyan overlay represents ROIs for the term “cognitive control” identified by meta-analysis. The demeaned regression coefficients (betas) of GLM4 for the main ROIs are displayed for each subject (dots and grey lines). The upper lines and stars represent significance (p < 0.025) of paired t-tests between conditions using the main contrasts ROIs These statistical comparisons are different from the main effect contrast used to define the ROIs and are shown for illustration purposes. They show that the regions do not respond to differences in stimulus mappings.

We further validated these results by analyzing the effects of stimulus discriminability on brain activity. Discriminability provides an additional, finer-grained manipulation of input-configuration salience. On each trial, we varied the difference in number of relevant elements between the two clouds (4%, 8%, and 12%), thereby increasing the saliency of the numerosity difference in the two stimuli. We included this discriminability measure (Figure 2, Figure 4a) as a parametric modulator in GLM1. As expected, and similar to the Sal Stim > Alt Stim contrast, activity in the SBN regions of interest (the bilateral TPJ, precuneus, dmPFC and left dlPFC) increased with the numerosity difference (Figure 4b, Supplementary Table 5). This provides additional support that activity in the SBN increases systematically with the saliency of input-configuration mappings.

**Figure 4.**
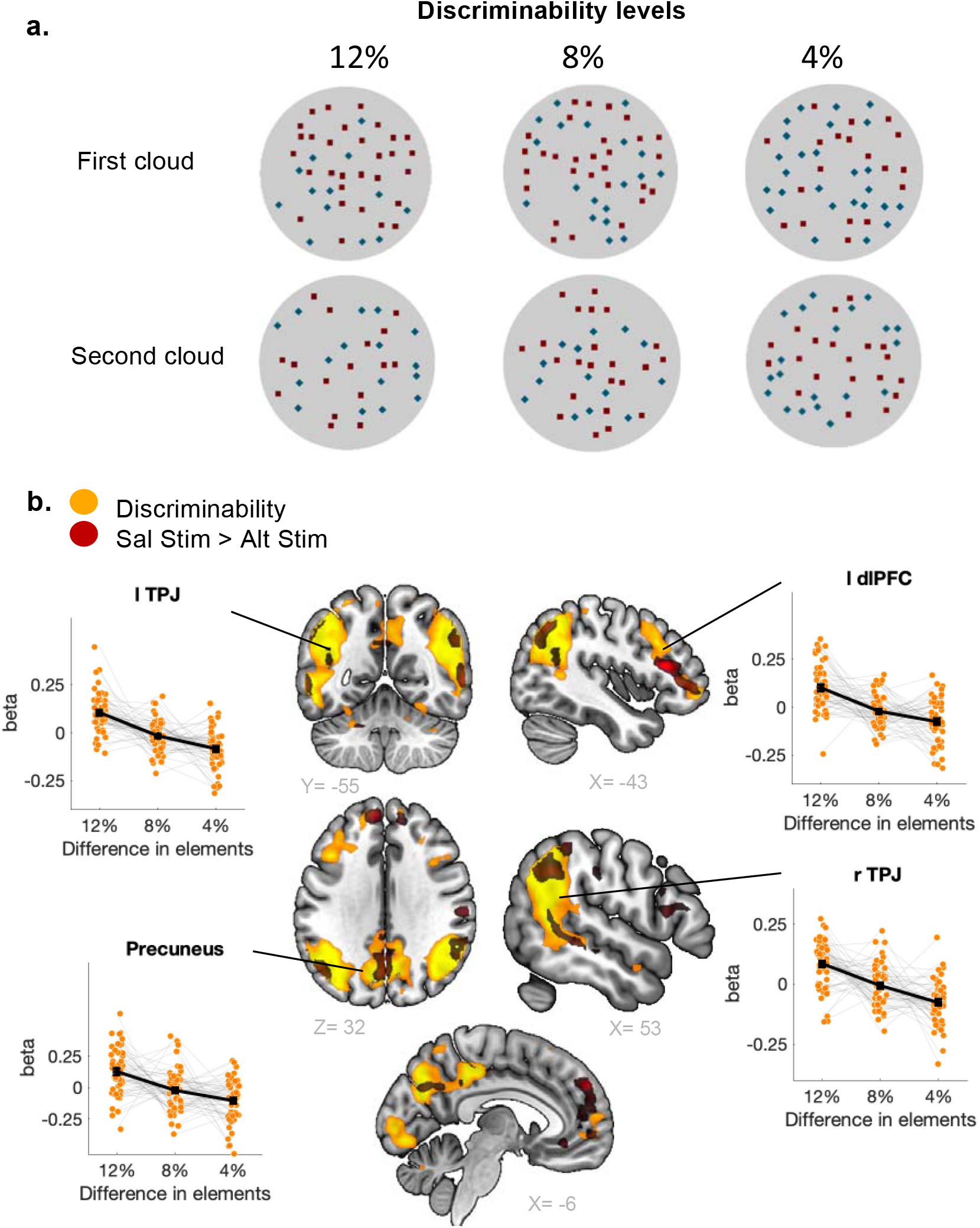
Stimulus discriminability, a second measure of configuration salience, leads to increased activity in SBN regions. (**a**) Example of the three discriminability levels for the salient stimulus condition (whole clouds). The differences in elements contained in the two clouds are 4%, 8%, and 12%. (**b**) Choice event, discriminability contrast. Clusters whose activation is significantly correlated with stimulus discriminability in the relevant stimulus condition (in yellow). The discriminability represents the difference in the proportion of behaviorally relevant stimulus (whole cloud for the salient condition, Alt Stim elements for the alternative condition) between the two clouds. The red overlay represents the contrast salient > alternative stimulus presented in Figure 3a. The demeaned regression coefficients (betas) of GLM5 for the main ROIs are displayed for each subject (orange dots and grey lines).

### Activity in the SBN does not reflect general task demands

We next confirmed that the results we observed did not reflect differences in general task demand, but rather selective recruitment of the SBN regions for salient input-configuration mappings. To do so, we analyzed the differences in brain activity between salient and alternative mappings in the response domain. These two response mappings were designed to match the difficulty of the salient and alternative stimulus mappings (confirmed by the matched RTs for the Alt Stim and Alt Resp conditions (Figure 2a)). We found that the Sal Resp > Alt Resp contrast did not show changes in the activity of most of the SBN regions (except for a cluster in the dmPFC). Instead, the posterior precuneus, left intraparietal lobule (IPL), and left posterior dlPFC were more activated in the alternative compared to the salient response condition (Figure 3b, Supplementary Table 2). These regions did not overlap with the “social” meta-analysis described above but instead overlapped with the meta-analysis obtained using the term “cognitive control” (Figure 3b, Supplementary Table 3). This result shows that the SBN is involved in the way stimuli are integrated but not in the way responses are selected during a choice process, and therefore specifically reflects information integration processes.

Despite our effort to match the difficulty of the alternative stimulus and response mappings (see Supplementary Methods), subjects were slightly more accurate in the alternative response (86%) than in the alternative stimulus condition (82%, Figure 2a, Supplementary Table 1). The difference in brain activation patterns between these two conditions could therefore be influenced by different levels of difficulty. To control for a potential difficulty confound reflected in this accuracy difference (but absent in the RT measures), we repeated all main analyses excluding 16 subjects with the highest difference in accuracy between the two alternative conditions (see Supplementary Methods and Supplementary Figure 2). The differences in RT and accuracy between the Alt Stim and Alt Resp conditions in this reduced dataset (38 subjects) were not significant (t-tests, accuracy p = 0.69, RT p = 0.37). The main results largely replicated in this control analysis, particularly the central result showing differential activity of the SBN regions for salient versus alternative input-configuration mappings (see Supplementary Figure 2c-d). The only difference that emerged was in the control analysis of the configuration-response mappings, where the dmPFC cluster activated in the Sal Resp-Alt Resp contrast was not significant. These analyses thus confirm that input-configuration and configuration-response mappings involve clearly separate brain networks, and that the involvement of the SBN in salient input-configuration mappings does not reflect task difficulty.

### Prefrontal regions show differential relation to the salience of stimulus versus response mappings

An area of specific interest in our analysis was the prefrontal cortex, since different studies have found parts of it to be involved in social cognition on the one hand^10,26,27,80^, and in cognitive control^46,47,81^ and choice execution^82,83^ on the other hand. This raises the question of whether prefrontal regions are recruited for salience-based stimulus integration (suggesting tight functional integration with the SBN) or for salience-based response selection (therefore being more involved in the cognitive control network that deals with general task demands) in our specific paradigm.

Our analyses suggest that the prefrontal cortex is involved in both these functions, but that different sub-areas respond to salience in the stimulus versus response domains. Overlaying the Sal Stim vs Alt Stim and Sal Resp vs Alt Resp contrasts (Figure 5a) revealed two clusters of the left dlPFC that responded to either stimulus salience or response salience, respectively, with increased activity for Sal Stim vs Alt Stim in the anterior part of the dlPFC (peak MNI coordinates −45, 26, 14, cluster extent 244, see Supplementary table 2), while a more posterior cluster was less activated in the Sal resp vs Alt resp conditions (peak MNI coordinates −42,5, 32, cluster extent 289, see Supplementary Table 2). These two clusters were mostly separated (22 voxels overlap). Thus, our results suggest that left dlPFC contains different (sub)areas that are functionally integrated with the SBN versus the cognitive control network; these regions are spatially close, with anterior regions governing input-configuration mappings and posterior regions more involved in configuration-response mappings.

**Figure 5.**
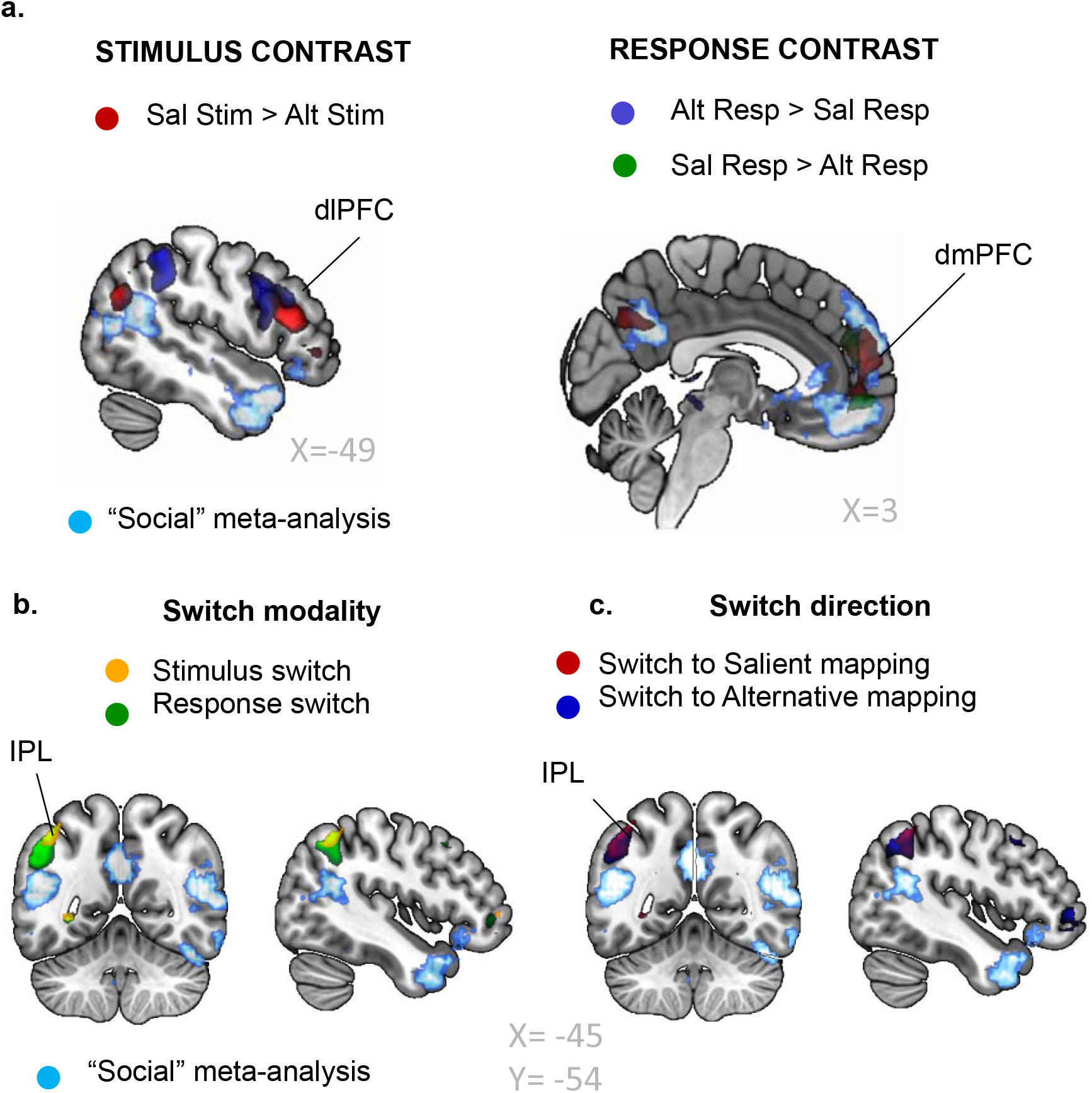
Gradients of activations in the prefrontal areas (a) and switch versus stay analysis (b-c). (**a**). Overlay of the main contrasts (Salient versus Alternative stimulus and response). The salient versus alternative contrasts of the choice event (Figure 3) are represented in red (stimulus contrast Sal Stim > Alt Stim), green (response contrast Sal Resp > Alt Resp), and blue (response contrast Alt Resp > Sal Resp). The cyan overlay represents ROIs for the term “social” identified by the NeuroSynth meta-analysis. (**b and c**) Contextual changes in switch versus stay trials do not recruit the SBN areas. (**b**) Choice event, switch vs stay trials: switch modality (stimulus or response). Clusters with significantly higher activity for the switch > stay trials following a stimulus switch (yellow) or a response switch (green) compared with stay trials. Switch trials are the first trials in a block (immediately following a switch in one of the two mappings), stay trials are trials 2 to 6 in a block. The cyan overlay represents ROIs for the term “social” identified by NeuroSynth meta-analysis. (**b**) Choice event, switch vs stay trials: switch direction (to alternative or to salient). Clusters with significantly higher activity for the switch > stay trials following an Alt ->Sal switch (red) or a Sal ->Alt switch (blue) compared with stay trials.

Activations in the dmPFC also appeared to follow a spatial gradient in response to stimulus and response salience (Figure 5a). A dorsal part of this region (peak MNI coordinates −9, 50, 35, cluster extent 403 see Supplementary Table 2) responded to the Sal stim vs Alt stim contrast, thus showing a similar pattern of activation than the other SBN regions. In contrast, a more ventral part of this region (peak MNI coordinates 6, 50, 8, cluster extent 188, see Supplementary Table 2) responded to salience in the response domain and was more active in the Sal resp > Alt resp contrast (Figure 5a – but note that this activation is not significant in the control analysis in which 16 subjects were excluded (Supplementary Figure 2)). These two clusters slightly overlap (41 voxels). The dmPFC therefore seems to be involved in salient mappings in both the stimulus and response domains, with a dorsal sub-area being responsible for input-configuration mappings and a ventral region potentially controlling configuration-response mappings.

### Salience-based activity in the SBN is not modulated by context switching

To evaluate the possible involvement of SBN regions (and in particular the rTPJ) in updating internal models of the environment after a contextual change^56^, we analyzed the brain activity linked to changes in contexts in our paradigm. Specifically, we conducted a switch versus stay trials analysis (GLM2 and GLM3), to examine the effect of changing contexts on the way information is integrated in different conditions. We compared the first trial in each block, where one of the mappings switched, to the rest of the trials. We included the type of switch (stimulus or response switch, Figure 5b, Supplementary table 7), or the direction of these switches (switch to salient or switch to alternative, Figure 5c, Supplementary table 7) in GLM2 and GLM3 respectively. We found that, for all types and directions of switches, the left inferior parietal lobule (IPL) was significantly activated for trials immediately following a switch, compared with trials occurring later in the blocks. This cluster did not overlap with the left TPJ from the “social” meta-analysis, suggesting that choices following a contextual switching in our task did not evoke increased activation in the TPJ.

For completeness, we additionally analyzed patterns of brain activity during the change-cue and feedback events, during which different updates of the model of the environment can occur. To do so, we included regressors for the change-cue (during which the model of the task is updated) and feedback (during which participants can learn if the model they used was accurate) events in GLM1. We found no significant differences in activations in the SBN regions for the contrasts Sal Stim > Alt Stim for the change-cue or feedback periods (Supplementary Figure 3). Note that while the change-cue analysis had lower power compared to the choice stage, we did not observe any activity clusters in the corresponding regions even with a relaxed threshold. This confirms that our results concerning activity in SBN regions reflects salience-based integration of stimuli into representations, rather than general contextual updating in the absence of stimuli.

## DISCUSSION

Regions thought to be crucially involved in social cognition (bilateral TPJ, dmPFC, left anterior dlPFC, and precuneus) are often also recruited in non-social tasks^33,40,41^. Previous theories have proposed that these findings reflect general functional mechanisms, but these mechanisms are usually still formulated in a rather task-specific manner (e.g., Theory of Mind^12^, Attention reorientation^33^ or updating of task context^56^). Here we propose an overarching functional mechanism that may unify the existing approaches and that may explain the recruitment of these regions across domains. We propose that these regions are involved in integrating multiple pieces of information into conceptually salient representations of world states that are used to guide behavior. We evaluated this proposal in the context of a new experimental task that allows us to dissociate salience of the input-configuration mappings from potential confounds such as task difficulty or contextual updating. We found that SBN areas were indeed specifically involved in mapping sensory information onto high-level configurations and were more active when the task required the subjects to focus on salient configurations. This salience-based effect was specific to stimulus integration (as opposed to motor salience) and did not reflect task difficulty or contextual updating.

### Social cognition re-expressed as salience-based stimulus integration in rTPJ

Our results refine theories of the rTPJ function^33,56^ that highlight the key role of this region in information processing across domains^60,61,64^. We propose a functional mechanism for this information integration, where multiple inputs are pieced together to form high-level representations. We show that SBN regions are more active when the task requires subjects to map inputs onto salient configurations that abstract away details and focus on the behaviorally relevant gist of the scene. This mechanism could explain the involvement of the rTPJ-dmPFC network in social computations such as TOM^41,54^. As shown by Heider and Simmel’s social attribution task, attributing social intentions, even to non-social (geometrical) interacting objects, is an automatic process, where different cues (relative movement of the different objects) are integrated to construct social representations^66^. Participants who are asked to describe the same geometrical object’s interactions in terms of single actors’ shapes and motions instead of social motives struggle to do so^84^. If thinking about other’s thoughts is a salient input-configuration mapping that requires the automatic grouping of various inputs into a behaviorally relevant abstract configuration, it is not surprising that the TOM network (dmPFC and rTPJ) is recruited by salient mappings in both social^12,16,54^ and non-social domains^40,43,44,85,86^.

Extending the TOM account of rTPJ function to more general social cognition, the “Nexus model”^41^ proposed that the rTPJ plays the role of an information processing hub that is specialized for creating a social context. Our proposal also encompasses this process and extends it, in considering social contexts to be particular occurrences of salient configurations. Processing social situations requires the integration of multiple inputs from the senses and memory that need to be brought together into a coherent social constellation that may match a template that is useful for action control. For example, to understand the movie proposal example from the introduction, we need to jointly consider the facial and gestural expressions of each character and perhaps their past interactions, to combine all this information and extract the behavioral gist of the situation. Yet, this seemingly complex process does not appear effortful but rather automatic in case of a salient, obvious interpretation. That salient social information may be integrated automatically by this network is also in line with the “social brain hypothesis”, according to which this network evolved to allow us to efficiently control our behavior in line with the omnipresent social information^3^. It would therefore not be surprising that a system specialized in processing behaviorally salient and overlearned types of information is recruited across many different types of social contexts.

Our proposal of a domain-general function of the SBN may appear at odds with reports of functional dissociations between subareas of this network (in particular the rTPJ-dmPFC network) that may be tuned to react in different ways to social versus non-social contexts^13,37,87^. This fine tuning of the type of information processed by subparts of the network does not rule out possible overarching functional mechanisms^53,88^, at the algorithmic level^52^. That is, different parts of a brain network may be specialized for processing different types of information but may still perform similar computations^51^. In this project, we sought to take a global approach and aimed at finding a mechanism that could explain why areas of the proposed SBN are reliably recruited across many different contexts. While our results are encouraging, it may be important to develop tasks that explicitly compare salience-based stimulus mappings between social and non-social stimuli, to reinforce the proposal put forth here. One way to do this may be to evaluate the brain activation of participants viewing social scenes while their task changes from focusing on the social meaning of the scene versus focusing on specific details. However, this will require careful matching of the sensory details between social and non-social contexts, which will not be trivial.

### Relation to other domain-general theories of rTPJ function

Previous theories have proposed various other domain-general mechanisms of rTPJ function, which are further refined by our proposal. The attention reorientation theory^33,55^ posits that the rTPJ is involved in reorienting attention to salient and behaviorally relevant stimuli. Here we confirm that salience^34^ and task relevance^35^ are indeed tightly linked to rTPJ activity, but in a way that transcends contexts that require reorientation. Building on the findings that a “circuit break” signal is inconsistent with the timing of activation of the rTPJ, Geng et al proposed a “contextual updating”^56^ function for this region. Recent work additionally supports the hypothesis that rTPJ is involved in updating beliefs in response to changes in the environment^89–91^ or impressions of others^92–94^. In our task, however, the rTPJ-dmPFC network appeared involved in stimulus information integration (Sal Stim >Alt Stim contrast) no matter whether this integration followed changes in task contexts. We did not find evidence that this differential activation was linked to an updating process or showed “circuit breaking” activity (differential activation in switch versus stay trials).

What activates this region during task changes may therefore not be the contextual update process per se, but rather the fact that the new context requires mappings of inputs to a more salient configuration. When individuals update their belief about a person (or the environment), they adapt the way they process the available information to draw conclusions that usually matches a template, and this template may change depending on the specific information. For example, when learning about the trustworthiness of others^92^, we may try to match the incoming information to templates of “trustworthy” or “untrustworthy” partners. It may be the integration of all pieces of incoming information to these salient templates, rather than a general updating process, that leads to activation in the TPJ. Further experiments should be conducted to differentiate these different proposals, for example by varying the type of contextual updates and comparing cases in which subjects update two equally salient contexts versus contexts of different salience.

### Dissociation of activity in the prefrontal cortex in stimulus and response domains

In addition to refining the rTPJ function, our results allow us to answer some unclarities about differential functional contributions of prefrontal cortex areas (dmPFC and dlPFC). The different regions of the SBN (in particular the TPJ and PFC) are often activated together, and, although differential activation has been reported in some tasks^93,95^, it remains unclear what precise mechanisms may be implemented by these different regions^96^. In the prefrontal cortex, two regions - the dmPFC and dlPFC - have both been associated with social computations, but also with cognitive control^46,47^ and behavioral flexibility in both social and non-social domains^48–50,97^. In our task, we found differential activations for these two regions. Both dorsal dmPFC and left anterior dlPFC were more activated in the salient compared to alternative stimulus mappings, showing their role in integration of task-relevant, salient, information, which has been previously associated with dlPFC activity^98^.

A ventral part of the dmPFC was more active in the salient compared to alternative response mapping, suggesting that the dmPFC area may be involved in salient mappings across domains, with sub-areas specifically dealing with salience in the stimulus versus response domains. By contrast, the left posterior dlPFC was more activated in the alternative compared to salient response condition, along with other regions belonging to the cognitive control network^46,47^. We noted a gradient of activation in the dlPFC, with the anterior part linked to stimulus integration rules and the posterior part being modulated by changes in response mappings, or action execution, in line with gradients of activity reported in the dlPFC during cognitive control tasks^46,82,99^. The dorsal dmPFC showed an activity pattern of activity that closely resembled the TPJ’s activations, suggesting that these regions support very similar functions. By contrast, the graded and differentiated pattern of activity in the dlPFC suggests that this area may play the role of a coordination hub, providing a link between stimulus integration and response execution networks.

### Relation to task demand and the default mode network

The default mode network (DMN) overlaps – although not fully – with the SBN^74^, and the regions we report in this analysis. This network is recruited when participants are at rest and deactivated during demanding tasks^45,100^. While this is a very common finding, there is considerable unclarity on what functional significance this activity may have during low-demand phases. In our study, we show that activity in SBN regions overlapping with this network was not related to lower task demand in general, but only increases for salient, less demanding stimulus integration (there was no such increase for the comparison of more versus less demanding motor selection). This suggests that the involvement of these areas indeed reflects a specific function in salience-based integration of stimuli into mental representations, rather than low task demand per se.

In turn, our results tentatively suggest a specific function that the DMN may be engaged in during the low-demand contexts used to study DMN activity. Since participants in these contexts are mostly free to perceive and/or think along lines that come to their mind, it is tempting to speculate that these regions may automatically map all incoming information (be it sensory or internal) onto salient high-level representations, without any constraints imposed by task instructions. This speculative proposal appears consistent with other proposals that the DMN may be involved in the construction of concepts^101^, in particular in the semantic domain^102^. Researchers who made the connection between resting state and social cognition propose a “social priming” mechanism^103^, which posits that rest periods prime the brain to process social information^104^, reinforcing the hypothesis that social computations may correspond to salient mappings that are automatically recruited^105^. Taken together, the function we propose here for the TPJ and other SBN regions also appears consistent with many empirical findings and speculations for the entirely separate literature on the DMN.

### What are salient configurations and how do they emerge?

In this project, we define salient configurations as automatically constructed representations of world states. They can be viewed as templates that allow us to interpret the world in an efficient manner, without wasting resources to focus on behaviorally irrelevant details. This definition raises the question of how these templates emerge. Are they innate, or do individuals learn them throughout their lives? Research on human development and social behavior seems to suggest that some of these templates emerge throughout human development. Theory of mind capacity for example – which may be closely related to the use of such templates - emerges during childhood^106^, and functional specialization of the TOM network increases with children’s age^107^. Lower-level representations, such as face recognition, seem to be present from a much earlier age^108^, and it remains debated to which extent this capacity is innate or learned^109^.

It is therefore possible that the high-level conceptual representations we focus on in this project rely on templates learned throughout development and experience. Regions belonging to the SBN, such as the TPJ, where various streams of sensory and mnemonic information converge, would then play the role of an integration hub where these templates and new information can be compared. Supporting this hypothesis, Vatansever and colleagues highlighted a possible role of the DMN in “autopilot” mode, where this network is activated when participants must select responses based on newly learned templates, while a different network supports the learning of these templates^110^. This hypothesis and our results open interesting research questions. How are different templates learned? How are cultural differences related to different templates, and are those reflected in the SBN activity?

### Implications for social neuroscience

Our new proposal and experimental findings have clear implications for the design of social neuroscience tasks, and the interpretation of their findings. If the SBN regions are activated when subjects construct conceptually salient representations, social neuroscience experiments should be carefully designed to consider differences in configurational salience. Typical tasks used in the social neuroscience literature may compare conditions with very different salience levels, so that the conclusions drawn from these tasks may be ambiguous. The activity reported as a difference between social and non-social computations may instead reflect differences in the salience of the representations constructed in the task and control conditions.

For instance, false belief versus false photograph tasks^12,111,112^ employ vignettes describing a situation involving either a person or an inanimate object. If the social information is more salient, and its integration is more automatic than processing information about an object, a confound is introduced. Similarly, other paradigms draw the subjects’ attention towards a character trait versus a grammatical construct of a sentence^113,114^. Here again, attributing character traits to a person may be more salient during this task than looking for specific grammatical attributes in a sentence.

To claim that the regions detected are specifically recruited by social computations, one would need to design control tasks that are fully matched to the experimental tasks, particularly in terms of salience of configurations and automaticity of information integration. At the most basic level, tasks that compare social and non-social situations should have identical visual or textual stimuli. However, as our paradigm shows, using the same stimulus across experimental and control conditions is not sufficient to rule out the effect of input-configuration mappings salience, as this effect depends on the task instructions and the mappings subjects must recruit in each experimental condition. To match configurational salience, one could control for the number of mental reasoning steps that are required for the task, as well as the average response time. Computational models could be helpful in this respect, in particular, to identify common or distinct information processing requirements of behavior in social versus non-social situations and to examine and compare their implementation at the neural level^51,52^.

### Stimulus salience in disorders of social behavior

In addition to the discussion of the functional mechanisms of the SBN regions, our results could provide new insight for understanding disorders such as the autism spectrum disorder (ASD).

Individuals with ASD often demonstrate altered theory of mind capacities, in failing to attribute thoughts to others that differ from their own^31,67^. In line with the assumption that this deficit may reflect impairments in the neural systems processing social information, there is evidence that the SBN’s activity differs between autistic and typically developing individuals^71–73,115^. However, some type of non-social information processing is also altered in individuals with ASD. For instance, they struggle with grouping information and creating categories or shortcuts, such as interpreting Gestalt representations^69,70^. Our proposed role of the SBN in salience-based integration into mental representations could therefore explain the impairments of individuals with ASD across the social (representing other minds) and non-social (representing Gestalts) domains. This could provide a better understanding of the neural causes of the impairments in this disorder, and potentially help to measure these as well as potentially remedy these dysfunctions by means of targeted interventions.

## METHODS

### Subjects

We recruited 57 healthy volunteers (33 females, 24 males; age 19-35, average 24.4) from the subject pool of the Department of Economics at the University of Zurich. This sample size was determined by conducting a power analysis using data from a behavioral pilot (N = 46, see Supplementary Methods) to observe a significant difference in behavior between our experimental conditions (using paired t-tests with a power of 0.8 and a significance level of 0.05). All subjects were right-handed, medication-free, and with normal or corrected-to-normal eyesight. All subjects provided written informed consent. The study was approved by the ethics committee of the Canton of Zurich.

We excluded runs with excessive motion during scanning from the fMRI analysis (head translation of more than 3 mm, or head rotation of more than 5 degrees). Three subjects were completely excluded from all analyses due to excessive motion occurring during multiple runs. We used the remaining 54 subjects for our analyses.

### Experimental design

We designed a new experimental task, systematically manipulating the salience of input-configuration and configuration-response mappings in a 2×2 within-subject design to disentangle behavioral and neural responses to mappings of different salience in stimulus and response domains. Subjects viewed the same stimuli in 4 experimental conditions, in which the rules for stimulus integration or response varied and performed a numerosity task (Figure 1a-c). On each trial, subjects observed two clouds of elements (Figure 1a-b) that were presented successively on a computer screen; each cloud had a different number of blue diamonds and red squares forming a circular patch. The subjects needed to compare the clouds and choose the one that had the highest number of elements.

To create different input-configuration mappings, we varied the perceptual task: In the salient stimulus (Sal Stim) condition, subjects had to select the cloud with the most elements, regardless of shapes and colors. In the alternative stimulus (Alt Stim) condition, subjects had to consider only one type of element (square or diamond) and ignore the second type, selecting the cloud that had the largest number of these specific elements. Thus, these two conditions imposed either the salient (whole cloud) or the alternative (specific elements) input-configuration mapping. We verified in a pilot phase that these two conditions indeed constituted a salient (and thus automatic and habitual) and an alternative (and thus more challenging and effortful) way of integrating the stimuli by comparing the accuracy and RT of subjects in the two conditions (see Supplementary Methods).

The specific element to attend in the Alt Stim condition was counterbalanced across subjects (squares: 26 subjects, diamonds: 31 subjects). We found no significant difference in accuracy or RT across the element type (t-tests, accuracy: p = 0.17, RT: p = 0.47).

To create matched difficulty configuration-response mappings, we varied the number of keypresses subjects had to provide to select each cloud. In the salient response (Sal Resp) condition, subjects had to press the response button once to select the first cloud and twice (similarly to a mouse double click) to select the second cloud. We confirmed with behavioral analysis that this constitutes an intuitive way of providing a task response (Supplementary methods and Figure 2a, low RT and high accuracy), as it uses the natural association between the sequential number of the cloud (1 or 2) and the number of keypresses needed to select these clouds (1 or 2 as well).

In the alternative response (Alt Resp) condition, subjects had to press once for the second cloud and twice for the first cloud. RT and accuracy (Figure 2a) confirmed that this reversed mapping constituted a non-intuitive, and more cognitively challenging way of providing a response to the task, as the natural association between numbers is altered. These two conditions manipulated the configuration-response mapping. As for the stimulus mappings, we validated that the two response mappings represented indeed a salient and an alternative response mapping in a pilot phase (see Supplementary Methods).

We manipulated the difficulty of each trial by varying the difference in the number of total and Alt Stim elements between the two clouds (4%, 8%, or 12% difference). This discriminability measure provides a second, more-fined grained measure of input-configuration salience that we used to validate our main hypothesis.

We additionally manipulated other parameters such as the total number of elements of the two clouds, or the proportion of Alt Stim elements relative to the total number of elements (See Supplementary Methods for details). Using the results of a pilot study, we constructed a set of 70 trials, which was counterbalanced for congruence of Alt Stim and Sal Stim conditions (See Supplementary Methods). The 70 trials were repeated 4 times (once per stimulus and response conditions), with an identical visual display, to construct the final set of 280 trials.

### Experimental task

The task was programmed using Psychophysics Toolbox Version 3^116^ and Matlab 2020a. The four conditions were presented in blocks of trials, each including 4 to 6 trials. After each block, one of the two conditions (stimulus or response) was randomly switched. At the beginning of each block, the subjects saw a cue presented for 2 s, which indicated which condition they would play in the coming block. The mapping (stimulus, indicated by a circle or a specific element, and response, indicated by “1-2” or “2-1”) that was being switched with respect to the previous block was displayed in blue.

The cue was then displayed in black for an additional 1.5 to 3.5 s (inter-stimulus interval, ISI). At the start of each trial, a fixation cross was displayed for 300 ms. The subjects then observed the two clouds of elements, shown for 500 ms each with an interval of 500 ms (intercloud interval, ICI). The subjects could reply as soon as the second cloud appeared and had 2.5 s to provide their response. Their response time (RT) was defined as the time elapsed between the onset of the second cloud and their first key press. A second key press was allowed for 400 ms after the first key press. The subjects then saw their response (number 1 or 2 indicating the chosen cloud) for 3 to 6 s (using a jittered interval sampled from a truncated gamma distribution).

After that, a feedback phase (1 s) indicated whether their answer was correct or incorrect. The task cue was then displayed (inter-trial interval, ITI) for 3 to 6 s (jittered, sampling from a truncated gamma distribution). If the subjects failed to reply within 2.5 s, a “time out” message was displayed, and the next trial started after the ITI (Figure 1d).

### Experimental session

Each session lasted for about 3 hours. After providing consent, the subjects completed a training phase outside the fMRI scanner. They performed 104 training trials. The first 16 trials were slow-paced (4 trials of each condition), to allow subjects to get used to the task. For these trials the events had the following duration: change cue = 4 s; ISI = 2 s; fixation = 1 s; clouds presentation and ICI = 800 ms; highlight answer = 3 s; feedback =1.5 s; ITI = 1.5 s. The next 72 trials were fast paced (ISI = 800 ms; highlight answer = 2700 ms; RT and ITI = 1200 ms, the other events had the same duration as described in Figure 1d). The last training phase outside the MRI scanner consisted of 16 real-paced trials, with the timing described in Figure 1d.

Inside the scanner, the subjects performed a short training of 8 randomly selected trials and then played the whole set of 280 trials (56 blocks). The trials were separated into 6 runs of 10 minutes, each including 46-47 trials.

### Subjects’ payoff

The subjects received a fixed payoff of 60 CHF (around 60 USD at the time of the experiment) for their participation. They additionally received money based on their average speed and accuracy throughout the session: For each percentage point of accuracy above 50% (chance level), they received 0.8 CHF; moreover, they received a total of 8 CHF if their average reaction time was below 500 ms, and a linearly decreasing proportion of this total if their reaction time was below 2000 ms. On average, this resulted in a total payoff of 94.33 CHF (min 83 CHF, max 102 CHF).

### Statistical analysis of behavioral data

We analyzed the accuracy and the reaction time (RT) of the subjects using (generalized) linear mixed-effect models (lme4 package in R). We used binomial regression to analyze the effect of the task design on subjects’ accuracy (glmer) and linear mixed effect models (lmer) to analyze the logarithm of the reaction time. As regressors, we included the stimulus and response conditions, the interaction between these two conditions, the position of the trial within a block (number of trials since the last switch) and the discriminability of the two clouds. Moreover, we included a random intercept as well as random slopes for each regressor for each subject in all analyses.

### fMRI data acquisition and pre-processing

The functional images were acquired using a Philips Achieva 3T whole-body scanner with a 32-channel Philips MR head coil. Each run was composed of 270 volumes and acquired with the following imaging parameters: 2334 ms repetition time (TR); 30 ms echo time (TE); 40 slices acquired in ascending order; 3 mm^2^ voxel size, 3 mm slice thickness; 0.5 mm gap; 90° flip angle. Five dummy image excitations were administered prior to the image acquisition stage. At the end of the session, a T1-weighted whole-brain structural image with 1×1×1 mm^3^ cubic voxel was recorded.

The preprocessing and analysis of the fMRI images were performed using SPM12 (Wellcome Trust Centre for Neuroimaging). All functional volumes were realigned to the first volume to correct for head movements. The 6 motion parameters (x, y, z translations and pitch, roll, and yaw rotations) were recorded and included at a later stage in the first-level GLM. Slice-timing correction was performed on the realigned images using the middle slice as a reference. The T1 (reference image) and the mean functional images (source image) were then coregistered and normalized to the T1 MNI template using the SPM12 segmentation procedure. The resulting images were finally smoothed with a 6mm FWHM Gaussian kernel.

### Peripheral measures

#### Physiological signals

Cardiac and respiratory signals were recorded throughout the experiment using MRI compatible electro-cardiogram and breathing belt. The physiological time series were transformed with the TAPAs toolbox^117^, using the RETROICOR method^118^ and included as a nuisance regressor in the first-level fMRI analysis.

#### Eye tracking

Eye motion was recorded using an MR-compatible infrared EYElink 1000 eye-tracker system (SR Research Ltd.). The position and size of the subject’s pupil were sampled at 500 Hz. The onsets and duration of saccades and blinks were recorded using the EYElink built-in function. Pupil size was linearly interpolated during eye blinks, z-transformed within each run, and averaged across micro-timesteps of 146 ms (1/16 of TR). The resulting variable as well as saccades and blinks were included as nuisance regressors in the first-level fMRI analysis, as we observed small differences between the experimental conditions (Supplementary figure 1c).

### First-level fMRI analysis

We conducted all first-level analyses using SPM12 (Wellcome Trust Centre for Neuroimaging). We constructed statistical general linear models (GLM) for each subject. We included a series of delta functions (time-locked to the events onsets) convolved with the canonical hemodynamic response functions (without its derivatives) in the design matrices.

In the main analysis (GLM1), we modeled 3 events as separate SPM conditions: choice, change cue, and feedback (Figure 1a). We added the stimulus and response experimental conditions (salient or alternative) as parametric modulators to the three regressors, as well as the modality that switched at the beginning of the block of trials (stimulus or response). In addition, we used the RT, the position of the trial in the block and their squared values, the discriminability measure (Figure 4b), the total number of elements N (Supplementary Figure 1) the congruence of the Sal Stim and Alt Stim conditions, and the correctness of the response as parametric modulators for the choice event. We additionally included 18 physiological, 5 eye-tracking regressors (see peripheral measures) and 6 motion parameters in the design matrices.

We constructed two additional GLMs to compare the brain activity for switch versus stay trials. These models were almost identical to the ones previously described. We did not include the parametric modulators coding for the trial positions in a block and instead included parametric modulators coding for switch versus stay trials. These parametric modulators were dummy variables taking the value 1 for trials immediately following a switch in the condition of interest and 0 for all other trials. In GLM2, we included response switch, and stimulus switch parametric modulators. In GLM3, we included the following parametric modulators: switching to a salient mapping and switching to an alternative mapping.

For illustration purposes, we constructed two additional GLMs (GLM4 and GLM5) to extract the regression weights corresponding to the 4 experimental mappings (Sal Stim/Sal Resp, Alt Stim/Sal Resp, Sal Stim/Alt Resp and Alt Stim/Alt Resp) and the 3 discriminability measures (4%, 8%, and 12%) from the ROIs extracted from the main analysis. Each experimental condition (mappings, or discriminability measure) was modeled as a separate condition for this analysis. We also included the trial position, RT, total number of elements, congruence, and discriminability (for the mappings analysis) as parametric modulators for each condition. The resulting regression weights (betas) were demeaned for each subject before display (Figure 3).

### Second-level fMRI analysis and correction for multiple comparisons

We conducted the second-level whole-brain univariate analyses using non-parametric testing (5000 permutations, no t-map smoothing) with SnPM (http://warwick.ac.uk/snpm). This method requires fewer assumptions than the classical parametric statistical approach^119^ and produces fewer false-positives^120^. We analyzed the contrasts of interest of the first-level level GLMs: Sal Stim > Alt Stim, Sal Resp > Alt Resp, discriminability, as well as the switch vs stay parametric modulators employing cluster-level correction with a threshold of FWE p < 0.05 using an initial cluster-forming threshold of p = 0.001 uncorrected. We added a dummy variable representing the type of Alt Stim elements (squares or diamonds), the average performance, average RT, and difference in accuracy and RT between Alt Stim and Sal Stim and between Alt Resp and Sal Resp for each subject as covariates in the second level analysis. The results are not significantly affected by inclusion or omission of these controls. In addition to the above-mentioned contrasts of interest, we analyzed the other parametric modulators included in our GLMs, as well as the change-cue and feedback events.

### Meta-analysis activation maps

We used an automated term-based meta-analysis with Neurosynth^79^ to download maps associated with the terms “social” and “cognitive control” (downloaded in November 2020). We used association test maps, displaying voxels that are more frequently reported in published articles that include the term compared with articles that do not (see FAQ on neurosynth.org). We used these maps to visualize overlaps between the activation clusters founds in our analysis and the clusters commonly found in the literature.

## Data availability

The data sets generated during and analyzed for the current study are publicly available in the OSF repository at: https://osf.io/7dgjw/

## Code availability

The code reproducing the analysis is publicly available in the OSF depository at: https://osf.io/7dgjw/

## Acknowledgements

This project has received funding from the European Research Council (ERC) under the European Union’s Horizon 2020 research and innovation programme (grant agreement No 725355, BRAINCODES). Claire Lugrin gratefully acknowledges the support of a Marlene-Porsche Foundation scholarship. We thank Christopher C. Hill for the helpful discussions.

## Author contributions

All authors designed the experiment and analyses. C.L. programmed the experiment. C.L. and A.K. conducted the experiment. C.L. performed the data analysis. All authors co-wrote the paper. C.C.R. supervised the project.

