## Supplementary material for "Salience-based information integration: An overarching function of the “social brain”?": Supplemetary material

### SUPPLEMENTARY METHODS

#### Trials selection

##### Trials construction

We first constructed a set of 112 trials of varying difficulty using the following procedure.

We systematically varied the total number of elements presented during a trial (65, 75, or 85), the difference in the number of elements between the two clouds (4%, 8%, or 12% difference), the difference in the number of the Alternative stimulus (Alt Stim) elements between the two clouds (4%, 8% or 12% difference), and the proportion of the Alt Stim elements relative to the total number of elements in the smallest cloud (0.35, 0.5 or 0.65).

Using these proportions, we first constructed all possible incongruent trials, in which the larger cloud had fewer Alt Stim elements than the smaller cloud (therefore leading to incongruent correct responses between the Sal Stim (salient stimulus) and Alt Stim conditions). We only included trials in which both clouds were composed of at least 11 squares and 11 diamonds. This yielded 56 trials.

We then created the congruent trials (for which the bigger cloud had more Alt Stim elements compared to the smaller cloud) matching these incongruent trials in all the above-mentioned proportions, generating a balanced design with equal numbers of congruent and incongruent trials.

The 112 congruent and incongruent trials were repeated 4 times (once per stimulus and response conditions), resulting in a set of 448 trials.

##### Behavioral pilots

We conducted two behavioral pilot sessions to verify that our experimental design elicited a differential response to salient and alternative mappings and to select the trials for the fMRI experiment.

Forty-six subjects participated in an online version of the experiment, and five subjects in an in-person version. All subjects faced the full trial sets of 448 trials and played the fast-paced version of the task (ISI = 800 ms; highlight answer = 2700 ms; RT and ITI = 1200 ms, the other events had the same duration as described in Figure 1). The online group participated using their personal computer with the browser Firefox and a version of the task programmed using JsPsych^1^. For the in-person group, the task was displayed using Psychophysics Toolbox Version 3 (PTB-3)^2^ running with Matlab 2020a. The pilot sessions lasted 1 hour 15 minutes. Subjects received on average 27 CHF for their participation (depending on their accuracy and speed).

These pilots allowed us to verify that the accuracy and RT were significantly affected by our experimental manipulations. The subjects were less accurate and slower in both the alternative stimulus and alternative response conditions.

##### fMRI trial selection

For the fMRI experiment, we selected a subset of 70 trials, to comply with fMRI time constraints and minimize the difficulty differences between the Alt Stim and Alt Resp conditions. We used the results from the two behavioral pilots to identify these trials. To keep a counterbalanced design including matching congruent and incongruent trials, we selected 35 trials, and their respective matching (congruent or incongruent) trials were also included in the final set. Each trial was then used 4 times, once per experimental condition, resulting in the final trial set of 280 trials.

Specifically, we used the online pilot data (N = 46 subjects, training set) to select 35 trials that had a minimal difference in performance between the two alternative conditions (Alt Stim and Alt Resp). The second in-person pilot (N = 5) was used as a validation set, to confirm that the trials we selected showed the expected behavior for a different group of subjects (that was not used in the trials selection process). We verified that the behavior of these subjects for the chosen 280 trials was matching the behavior of the 46 subjects used for the trial selection procedure and that there were no significant differences in accuracy and RT between the two alternative conditions for the two separate groups of subjects.

##### fMRI sample size computation

We used the size of the effects of the alternative versus salient conditions on the accuracy and RT of the 46 online subjects (restricted to the 280 selected trials) to determine the sample size needed for our fMRI study. We computed the sample size needed to observe a significant difference in accuracy and RT between the salient and alternative for both the response and stimulus conditions, using paired t-tests with a power of 0.8 and a significance level of 0.05.

#### Control fMRI analysis – difficulty

To control for the results of our fMRI analysis and rule out a possible detection of activity related to the general difficulty of our task, we conducted a control analysis in which we excluded 16 subjects who had a strong difference in accuracy between the alternative stimulus and the alternative response condition. We excluded these subjects to match the accuracy and RT at the group level for the alternative stimulus and response. We iteratively removed the subjects that had the highest Alt Stim - Alt Resp accuracy difference, until the derivative of the group level Alt Stim - Alt Resp difference became negative (Supplementary figure 1a). The Alt Stim - Alt Resp accuracy and RT difference of the new subjects’ group was nonsignificant (Supplementary figure 1b), indicating a matched difficulty between these two conditions. We then replicated the fMRI 1^st^ and 2^nd^ level analysis as described in the methods section using the subgroup of 38 subjects.

### SUPPLEMENTARY TABLES

**Supplementary Table 1.** **Statistical analysis.** Fixed effects coefficient estimates, standard errors, t-values, and p-values of the choice and RT regressions mixed-effects models using subjects as random effects. **a**. The choice data was analyzed using a binomial probit model. **b**. The log reaction time was analyzed using a linear model

| **a.** *Correct response ~ stimulus type * response type + trial position + discriminability + (1 + stimulus type * response type + trial position + discriminability \| subject)* | | | | |
| --- | --- | --- | --- | --- |
|  | Estimate | Std. Error | z value | p value |
| Intercept | -12.65 | 0.788 | -16.05 | <0.001 |
| stimulus type | -0.828 | 0.093 | -8.82 | <0.001 |
| response type | -0.252 | 0.079 | -3.19 | <0.001 |
| trial position | 0.048 | 0.015 | 3.07 | 0.002 |
| discriminability | 27.14 | 1.467 | 18.50 | <0.001 |
| Stimulus type * Resp type | 0.078 | 0.104 | 0.74 | 0.455 |
| **b.** *log(RT) ~ stimulus type * response type + trial position + (1 + stimulus type * response type + trial position \| subject)* \| \| \| \| \|; \| Estimate \| Std. Error \| df \| t value \| p value; Intercept \| 0.571 \| 0.122 \| 53.12 \| 4.69 \| <0.001; stimulus type \| 0.140 \| 0.010 \| 59.39 \| 13.36 \| <0.001; response type \| 0.126 \| 0.011 \| 57.85 \| 11.83 \| <0.001; trial position \| -0.008 \| 0.002 \| 55.03 \| -4.38 \| <0.001; discriminability \| -2.002 \| 0.195 \| 52.37 \| -10.22 \| <0.001; Stimulus type * Resp type \| -0.046 \| 0.010 \| 167.13 \| -4.48 \| <0.001 | | | | |

**Supplementary Table 2. Salient > Alternative contrasts.** Corresponds to Figure 3. BOLD contrast showing significant clusters (FWE < 0.05) for salient > alternative stimulus and response mappings. T-statistics and cluster-forming threshold are calculated using SnPM13 (see Methods): t > 3.25, threshold = 57 voxels. MNI coordinates show peak activations for each cluster.

| **GLM1, Contrast Name: Salient Stimulus > Alternative Stimulus** | | |  |  | MNI Coordinates | | |
| --- | --- | --- | --- | --- | --- | --- | --- |
|  | Region | Region Label | Extent | t-value | x | y | z |
| Positive | R TPJ | R SupraMarginal Gyrus | 230 | -7.428 | 63 | -28 | 44 |
| Sal Stim > Alt Stim |  | R Middle Temporal Gyrus | 156 | -5.215 | 60 | -52 | 5 |
|  | dmPFC | L Superior Medial Gyrus | 403 | -5.315 | -9 | 50 | 35 |
|  | L TPJ | L Angular Gyrus | 222 | -5.130 | -42 | -64 | 50 |
|  |  | L Middle Temporal Gyrus | 119 | -5.231 | -63 | -49 | 5 |
|  | Precuneus | L Precuneus | 97 | -4.722 | 0 | -67 | 35 |
|  | L dlPFC | L Middle Frontal Gyrus | 59 | -4.673 | -36 | 14 | 50 |
|  |  | L IFG (p. Triangularis) | 244 | -6.450 | -45 | 26 | 14 |
|  | R Insula | R Insula Lobe | 76 | -4.591 | 42 | -7 | 5 |
| Positive | L Occipital | L Middle Occipital Gyrus | 746 | 10.744 | -33 | -85 | 17 |
| Alt Stim > Sal Stim | R Occipital | R Middle Occipital Gyrus | 675 | 9.217 | 36 | -79 | 23 |
| **GLM1, Contrast Name: Salient Response > Alternative Response** | | |  |  | MNI Coordinates | | |
|  | Region | Region Label | Extent | t-value | x | y | z |
| Positive | dmPFC | R Superior Medial Gyrus | 188 | -4.518 | 6 | 50 | 8 |
| Sal Resp > Alt Resp |  | R Superior Frontal Gyrus | 98 | -4.360 | 24 | 29 | 50 |
|  | Heschls Gyrus | R Heschls Gyrus | 68 | -4.631 | 39 | -25 | 14 |
| Negative | L dlPFC | L Middle Frontal Gyrus | 147 | 6.872 | -36 | 2 | 56 |
| Alt Resp > Sal Resp |  | L IFG (p. Opercularis) | 289 | 5.474 | -42 | 5 | 32 |
|  | L IPL |  | 517 | 6.608 | -36 | -43 | 41 |
|  | Precuneus | L Precuneus | 87 | 6.263 | -6 | -67 | 47 |
|  | STN |  | 137 | 5.362 | -12 | -16 | -10 |

**Supplementary Table 3. Extents of overlap with the “social” and “cognitive control” meta-analysis.**

| **GLM1, Contrast Name: Salient Stimulus > Alternative Stimulus** | |
| --- | --- |
| Region | Overlap with “Social” meta-analysis |
| dmPFC | 289 voxels |
| R TPJ | 55 + 16 voxels |
| L TPJ | 52 voxels |
| Precuneus | 60 voxels |
| **GLM1, Contrast Name: Alternative Response > Salient Response** | |
| Region | Overlap with “Cognitive control” meta-analysis |
| dlPFC | 121 voxels |
| L IPL | 18 voxels |

**Supplementary Table 4, Difficulty matched control analysis: Salient > Alternative contrasts.** Corresponds to Supplementary figure 2a. BOLD contrast showing significant clusters (FWE < 0.05) for salient > alternative stimulus and response mappings using the reduced data set of 38 subjects. T-statistics and cluster-forming threshold are calculated using SnPM13 (see Methods): t > 3.38, threshold = 57 voxels. MNI coordinates show peak activations for each cluster.

| **GLM1 CONTROL, Contrast Name: Salient Stimulus > Alternative Stimulus** | | |  |  | MNI Coordinates | | |
| --- | --- | --- | --- | --- | --- | --- | --- |
|  | Region | Region Label | Extent | t-value | x | y | z |
| Positive | R TPJ | R SupraMarginal Gyrus | 187 | -6.982 | 60 | -40 | 47 |
| Sal Stim > Alt Stim |  | R Middle Temporal Gyrus | 111 | -5.704 | 63 | -52 | 8 |
|  | dmPFC | L Mid Orbital Gyrus | 263 | -4.751 | -3 | 59 | -4 |
|  | L TPJ | L Middle Temporal Gyrus | 266 | -6.113 | -63 | -49 | 2 |
|  | Precuneus | L PCC | 145 | -5.119 | 0 | -58 | 32 |
|  | L dlPFC | L Middle Orbital Gyrus | 72 | -4.984 | -36 | 50 | -4 |
|  |  | L IFG (p. Triangularis) | 61 | -4.898 | -45 | 26 | 17 |
|  | R Insula | R Insula Lobe | 78 | -5.396 | 39 | 5 | 5 |
| Positive | L Occipital | L Middle Occipital Gyrus | 453 | 8.157 | -33 | -88 | 20 |
| Alt Stim > Sal Stim | R Occipital | R Middle Occipital Gyrus | 340 | 6.563 | 36 | -79 | 23 |
| **GLM1 CONTROL, Contrast Name: Salient Response > Alternative Response** | | |  |  | MNI Coordinates | | |
|  | Region | Region Label | Extent | t-value | x | y | z |
| Negative | L dlPFC | L IFG (p. Triangularis) | 922 | 7.517 | -48 | 8 | 32 |
| Alt Resp > Sal Resp | L IPL + precuneus | L Inferior Parietal Lobule | 836 | 9.195 | -54 | -40 | 47 |
|  | STN |  | 260 | 6.928 | -9 | -22 | -10 |

**Supplementary Table 5. Clouds discriminability contrast.** Corresponds to Figure 4. BOLD contrast showing significant clusters (FWE < 0.05) correlating with the discriminability parametric modulator. T-statistics and cluster-forming threshold are calculated using SnPM13 (see Methods): t > 3.25, threshold = 57 voxels. MNI coordinates show peak activations for each cluster.

| **GLM1, Contrast Name: Stimulus Discriminability (pmod)** | | |  |  | MNI Coordinates | | |
| --- | --- | --- | --- | --- | --- | --- | --- |
|  | Region | Region Label | Extent | t-value | x | y | z |
| Positive | R TPJ | R Angular Gyrus | 1471 | 8.979 | 51 | -58 | 32 |
| Increase with |  | R Superior Temporal Gyrus | 1471 | 3.540 | 51 | -46 | 17 |
| discriminability | L TPJ | L Angular Gyrus | 3240 | 8.397 | -48 | -61 | 41 |
|  | Prefrontal cortex: dmPFC and bilateral dlPFC | L Middle Frontal Gyrus | 1778 | 6.505 | -36 | 11 | 53 |
|  |  | L Middle Orbital Gyrus | 171 | 6.175 | -39 | 56 | -1 |
|  |  | R Middle Orbital Gyrus | 58 | 5.019 | 39 | 41 | -10 |
|  | Precuneus | R Precuneus | 3240 | 3.594 | 9 | -58 | 44 |
|  | ParaHippocampal | L ParaHippocampal Gyrus | 146 | 5.401 | -27 | -19 | -19 |
|  | Cerebellum | Cerebellum | 117 | 5.011 | -15 | -70 | -31 |

**Supplementary Table 6. Change cue and feedback events contrast.** Corresponds to Supplementary Figure 3. BOLD contrast showing significant clusters (FWE < 0.05) for Response > Stimulus switch during a change cue event. T-statistics and cluster-forming threshold are calculated using SnPM13 (see Methods): t > 3.25, threshold = 57 voxels. MNI coordinates show peak activations for each cluster.

| **GLM1, Contrast Name: Change cue: Switch stimulus > Switch response** | | | | MNI Coordinates | | |
| --- | --- | --- | --- | --- | --- | --- |
|  | Region Label | Extent | t-value | x | y | z |
| Positive | L Inferior Temporal Gyrus | 177 | -6.0265 | -48 | -52 | -13 |
| Switch Stim > Switch Resp | L IFG (p. Opercularis) | 108 | -5.6523 | -39 | 2 | 29 |
|  | L IFG (p. Triangularis) | 79 | -5.5928 | -51 | 35 | 14 |
| Negative | L Inferior Parietal Lobule | 191 | 6.49235 | -57 | -52 | 41 |
| Switch Resp > Switch Stim | ACC | 442 | 6.42932 | 18 | 29 | 29 |
|  | R SupraMarginal Gyrus | 132 | 4.92867 | 60 | -43 | 44 |
| **GLM1, Contrast Name: Feedback: Alternative stimulus > Salient Stimulus** | | | | MNI Coordinates | | |
|  | Region Label | Extent | t-value | x | y | z |
| Positive | Cerebellar Vermis (8) | 431 | 7.31415 | 0 | -76 | -28 |
| Alt Stim > Sal Stim | L Putamen | 128 | 6.95683 | -18 | 11 | -4 |
|  | R Putamen | 204 | 6.24425 | 24 | 14 | -4 |
|  | L Cerebelum (Crus 1) | 160 | 5.25061 | -36 | -64 | -28 |
|  | L Precentral Gyrus | 73 | 4.98877 | -33 | -19 | 62 |
|  | R Thalamus | 57 | 4.70959 | 6 | -19 | 8 |
|  | L Postcentral Gyrus | 173 | 4.67488 | -45 | -28 | 47 |
|  | L Fusiform Gyrus | 62 | 4.4261 | -39 | -61 | -4 |

**Supplementary Table 7. Switch versus stay contrasts.** Corresponds to Figure 5b. BOLD contrast showing significant clusters (FWE < 0.05) for Switch > Stay trials, for the different switching modalities. T-statistics and cluster-forming threshold are calculated using SnPM13 (see Methods): t > 3.25, threshold = 57 voxels. MNI coordinates show peak activations for each cluster.

| **GLM2, Contrast Name: Switch vs stay trials: Switch stimulus** | | | | MNI Coordinates | | |
| --- | --- | --- | --- | --- | --- | --- |
|  | Region Label | Extent | t-value | x | y | z |
| Switch Stimulus Positive | L Angular Gyrus | 70 | 6.138 | -42 | -61 | 50 |
| **GLM2, Contrast Name: Switch vs stay trials: Switch Response** | | | | MNI Coordinates | | |
|  | Region Label | Extent | t-value | x | y | z |
| Switch Response Positive | L Inferior Parietal Lobule | 185 | 7.186 | -51 | -55 | 47 |
|  | R ParaHippocampal Gyrus | 66 | 5.557 | 36 | -46 | -1 |
| **GLM3, Contrast Name: Switch vs stay trials: Switch to alternative** | | | | MNI Coordinates | | |
|  | Region Label | Extent | t-value | x | y | z |
| Switch to alternative positive | L Inferior Parietal Lobule | 182 | 6.970 | -45 | -58 | 50 |
| **GLM3, Contrast Name: Switch vs stay trials: Switch to salient** | | | | MNI Coordinates | | |
|  | Region Label | Extent | t-value | x | y | z |
| Switch to salient positive | L Inferior Parietal Lobule | 121 | 5.445 | -42 | -55 | 56 |

### SUPPLEMENTARY FIGURES

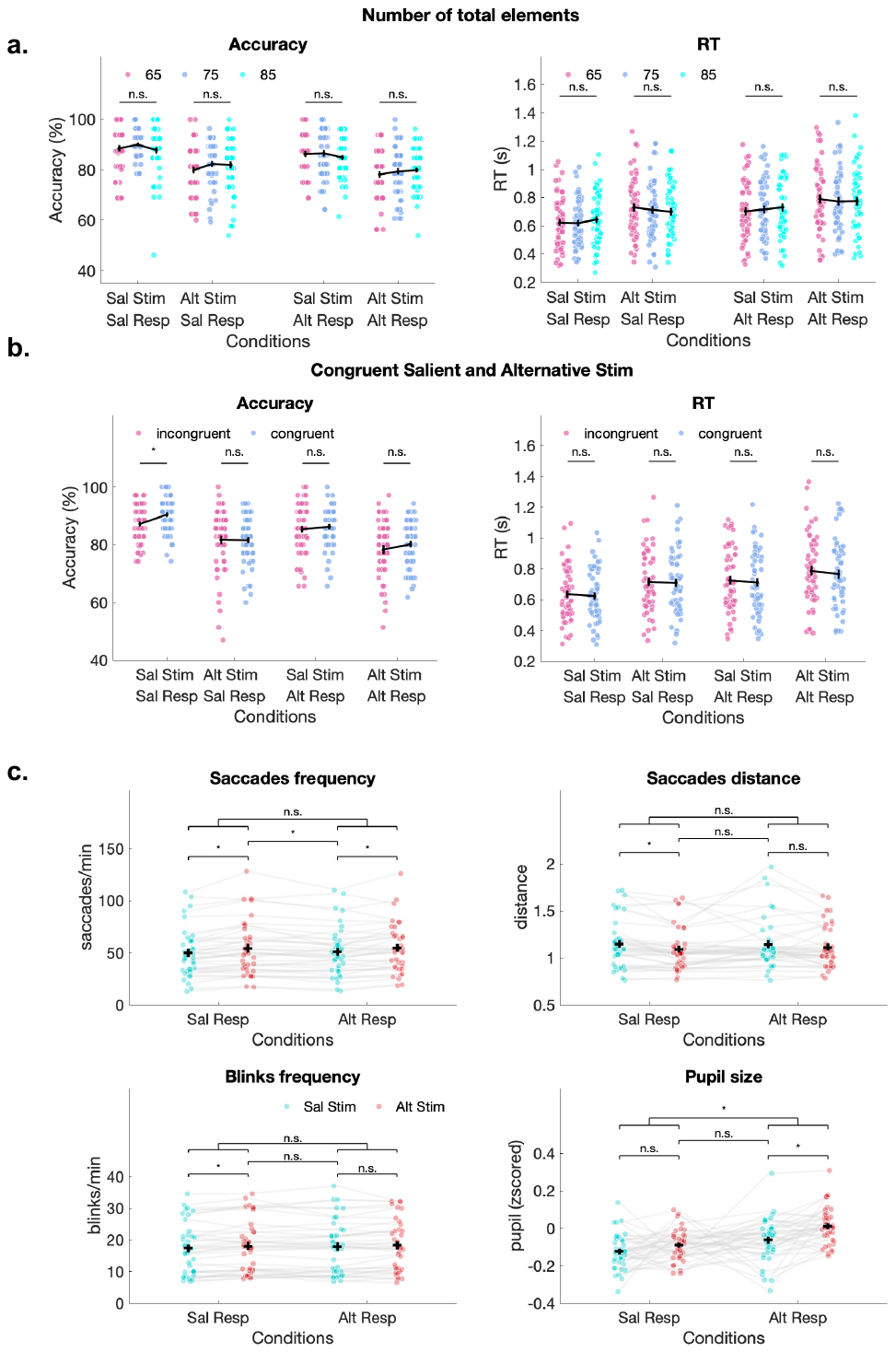

*Supplementary Figure 1. Behavioral and eye tracker controls. Corresponds to Figure 2. (a) Effect of the total number of elements on accuracy and RT. Each dot represents one subject, errors bars represent s.e.m. The horizontal lines and stars represent significance (p < 0.05) of linear regressions between accuracy (RT) and the total number of elements. (b) Effect of the congruence between the salient and alternative stimulus conditions on RT and accuracy. Each dot represents one subject, errors bars represent s.e.m. The horizontal lines and stars represent significance (p < 0.0125) of paired t-tests between conditions. (c) Eye tracking control of the number of saccades, distance of these saccades, numbers of blinks and pupil size in the 4 different conditions. Each dot and grey line represent one subject, errors bars represent s.e.m. The horizontal lines and stars represent significance (p < 0.0125) of paired t-tests between conditions.*

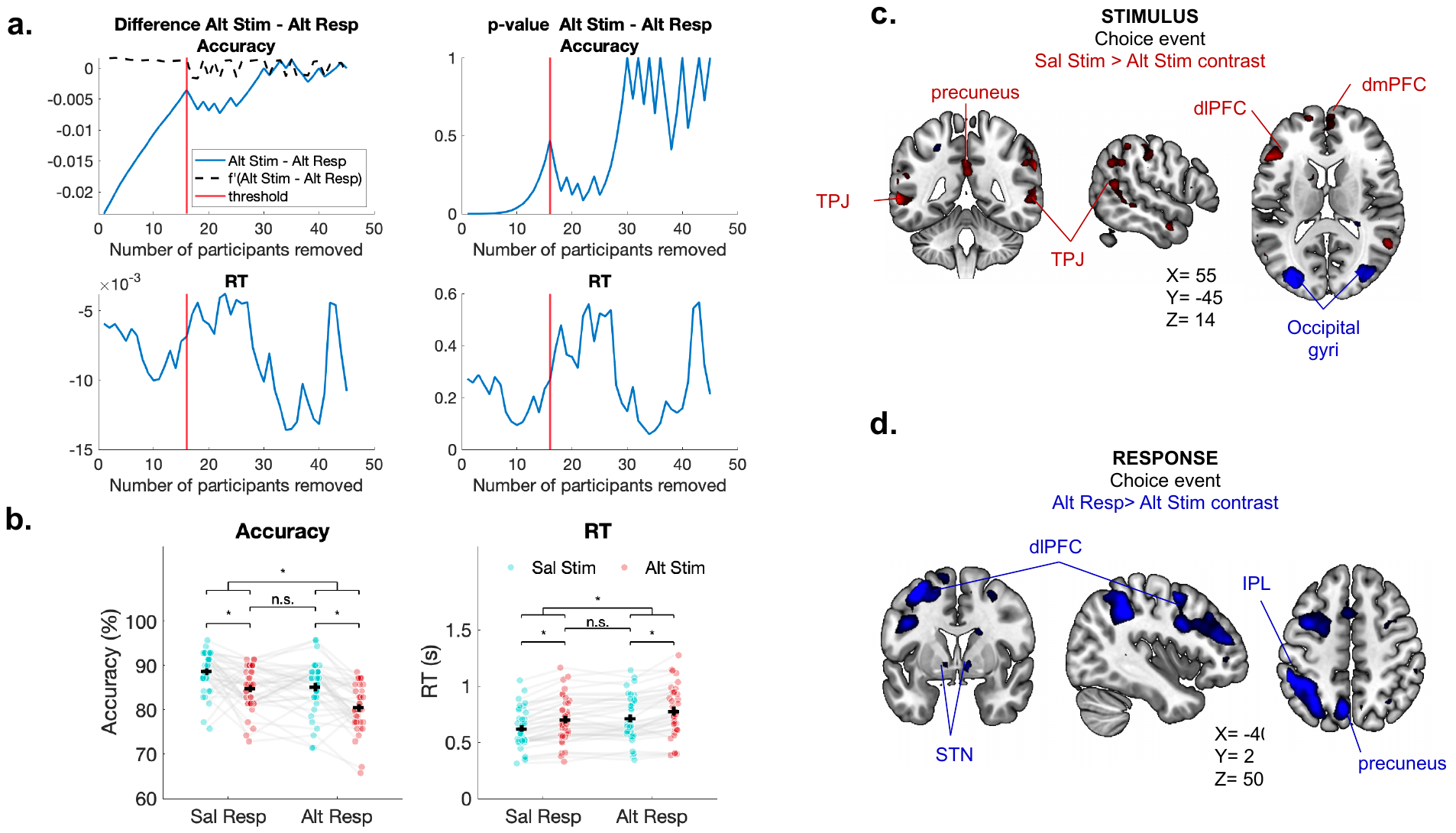

*Supplementary Figure 2. Procedure and results from matched difficulty control analysis. Corresponds to
Figure 3. (a) Subjects’ selection procedure: subjects are iteratively removed based on their absolute accuracy difference between the alternative stimulus and alternative response conditions. The threshold to stop removal is defined as the number of subjects for which the derivative of the Alt Stim - Alt Resp difference becomes negative. The effect of subject removal on RT differences, as well as the p-value of the Alt Stim - Alt Resp difference for accuracy and RT are also displayed. (b) Average accuracy and reaction time difference between the salient and alternative mappings in the stimulus or response modalities after 16 subjects are removed from the analysis. Each dot represents one subject, errors bars represent s.e.m. (c) Choice event, salient versus alternative stimulus contrast after 16 subjects are excluded from the analysis. Clusters with significantly higher activity for Salient > Alternative stimulus conditions (in red) or higher activity for Alternative > Salient stimulus conditions (in blue). (d) Choice event, salient versus alternative response contrast after 16 subjects are excluded from the analysis. Clusters with significantly higher activity for Alternative > Salient response conditions (in blue).*

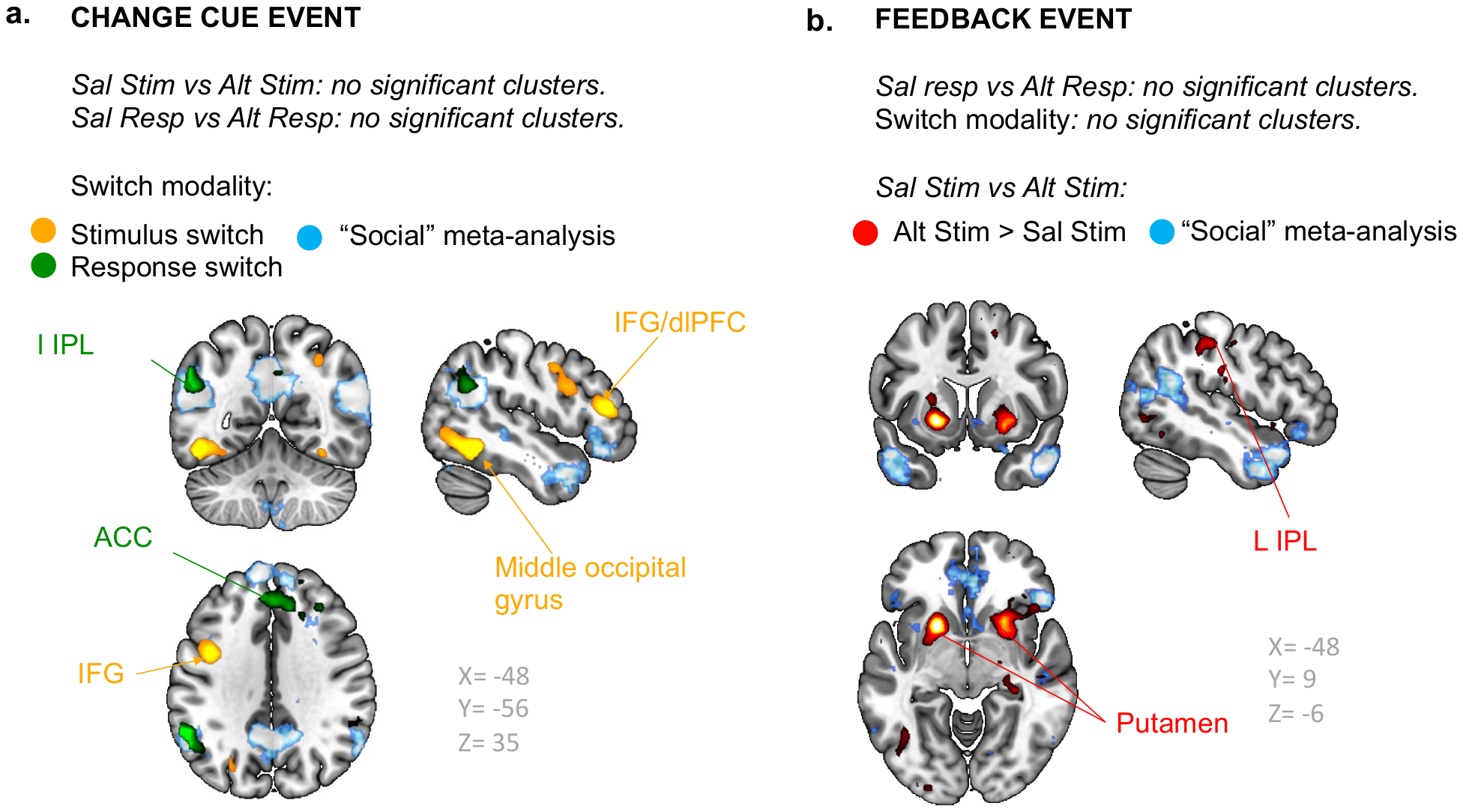

*Supplementary Figure 3. Control analysis, change cue, and feedback events. (a) Change cue event, stimulus versus response switch. Clusters with significantly higher activity for stimulus > response switches during the change cue event (in yellow) or higher activity for response > stimulus switches (in green). The cyan overlay represents ROIs for the term “social” identified by the NeuroSynth meta-analysis. We found no significant clusters for the salient>alternative contrasts for either the stimulus or response domains. (b) Feedback event, alternative > salient stimulus contrast. Clusters with significantly higher activity for the Alt Stim> Sal Stim trials during the feedback phase. We found no significant clusters in the salient versus alternative response mappings analysis.*
